# HERI-1 is a Chromodomain Protein that Negatively Regulates Transgenerational Epigenetic Inheritance

**DOI:** 10.1101/384792

**Authors:** Roberto Perales, Daniel Pagano, Gang Wan, Brandon Fields, Arneet L. Saltzman, Scott G. Kennedy

**Affiliations:** Department of Genetics, Harvard Medical School, Boston MA, 02115; Laboratory of Genetics, University of Wisconsin-Madison, Madison WI, 53706; Department of Cell and Systems Biology, University of Toronto, Toronto ON M5S 3G5

**Author notes:** Corresponding author, Department of Genetics, Harvard Medical School, 77 Avenue Louis Pasteur, New Research Building 266, Boston, MA. 02215.

## Abstract

Transgenerational epigenetic inheritance (TEI) is the inheritance of epigenetic information for two or more generations. In most cases, TEI is limited to 2-3 generations. This short-term nature of TEI could be set by innate biochemical limitations to TEI or by genetically encoded systems that actively limit TEI. dsRNA-mediated gene silencing (RNAi) can be inherited in *C. elegans* (termed RNAi inheritance or RNA-directed TEI). To identify systems that might actively limit RNA-directed TEI, we conducted a forward genetic screen for factors whose mutation enhanced RNAi inheritance. This screen identified the gene <u>h</u>eritable <u>e</u>nhancer of <u>R</u>NA<u>i</u> (*heri-1*), whose mutation causes RNAi inheritance to last longer (>20 generations) than normal. *heri-1* encodes a protein with a chromodomain and a kinase-homology domain that is expressed in germ cells and localizes to nuclei. In *C. elegans*, a nuclear branch of the RNAi pathway (nuclear RNAi or NRDE pathway) is required for RNAi inheritance. We find that this NRDE pathway is hyper-responsive to RNAi in *heri-1* mutant animals, suggesting that a normal function of HERI-1 is to limit nuclear RNAi and that limiting nuclear RNAi may be the mechanism by which HERI-1 limits RNAi inheritance. Interestingly, we find that HERI-1 binds to genes targeted by RNAi, suggesting that HERI-1 may have a direct role in limiting nuclear RNAi and, therefore, RNAi inheritance. Surprisingly, recruitment of the negative regulator HERI-1 to genes depends upon that same NRDE factors that drive co-transcriptional gene silencing during RNAi inheritance. We therefore speculate that the generational perdurance of RNAi inheritance is set by competing pro- and anti-silencing outputs of the NRDE nuclear RNAi machinery.

## Introduction

The inheritance of epigenetic information for two or more generations is referred to as transgenerational epigenetic inheritance (TEI) (Heard and Martienssen 2014). Many examples of TEI have now been documented including, but not limited to, paramutation in plants (Arteaga-Vazquez and Chandler 2010), protein-based inheritance in yeast (Shorter and Lindquist 2005), dsRNA-mediated gene silencing (RNAi) in *C. elegans* (Vastenhouw et al. 2006; Buckley et al. 2012a; Ashe et al. 2012), and the inheritance of acquired traits in mice (Carone et al. 2010; Walker and Gore 2011; Radford et al. 2012; Padmanabhan et al. 2013; Castel and Martienssen 2013; Somer and Thummel 2014; Holoch and Moazed 2015; R. Martienssen and Moazed 2015; Rankin 2015). In many cases, TEI is limited to a small number (*e.g*. 2-3) of generations (Heard and Martienssen 2014; D’Urso and Brickner 2014). In other cases, such as piRNA-mediated gene silencing in *C. elegans* and paramutation in plants, TEI can be long-lasting (>10 generations). Molecular mechanisms that set the generational duration of TEI are largely a mystery.

Recently, small non-coding RNAs have emerged as important mediators of epigenetic inheritance in eukaryotes. For example, in plants, the RNA-dependent RNA Polymerase (RdRP) mop1 produces small interfering (si)RNAs thought to mediate paramutation (Alleman et al. 2006). In the yeast *S. pombe*, siRNAs help direct and maintain stable, and in some cases heritable, heterochromatic states at pericentromeres (Volpe et al. 2002; R. A. Martienssen, Zaratiegui, and Goto 2005; Ragunathan, Jih, and Moazed 2015). In *Drosophila*, maternally inherited PIWI-interacting RNAs (piRNAs) direct heritable silencing of transposable elements (Le Thomas et al. 2014). Finally, in mice the effects of stress and metabolic disease are reported to pass from parent to progeny and miRNAs and short tRNA fragments have been implicated in mediating this inheritance (Carone et al. 2010; Gapp et al. 2014; Sharma et al. 2016). Thus, small regulatory RNAs are major mediators of TEI in eukaryotes (termed RNA-directed TEI). Small non-coding RNAs are also linked to TEI in *C. elegans*. For instance, piRNAs can trigger transgenerational gene silencing via a process termed RNA epigenetic (RNAe) (Ashe et al. 2012; Shirayama et al. 2012). Additionally, dsRNA-mediated gene silencing (RNA interference or RNAi) is heritable in *C. elegans*: the progeny of animals treated with dsRNAs retain the ability to silence RNAi-targeted genes for many generations, even after the removal of dsRNA triggers (termed RNAi inheritance) (Vastenhouw et al. 2006; Rosa M. Alcazar, Lin, and Fire 2008). During RNAi inheritance in *C. elegans*, double stranded RNAs (dsRNA) are processed by Dicer into “primary” small interfering RNAs (siRNAs), which are bound by Argonaute (AGO) proteins to regulate complementary cellular RNAs (Meister 2013). AGO-bound primary siRNAs recruit, by an unknown mechanism, RdRPs to homologous mRNA templates to produce amplified pools of “secondary siRNAs” (Sijen et al. 2001). Repeated RdRP-based siRNA amplification in germ cells each generation is likely responsible for helping to promote RNAi inheritance in *C. elegans* (Buckley et al. 2012b; Ashe et al. 2012; Sapetschnig et al. 2015).

Forward genetic screens have identified factors that are required for RNAi inheritance in *C. elegans* (Buckley et al. 2012b; Ashe et al. 2012; Spracklin et al. 2017; Wan et al. 2018). The factors fall into two general categories. The first category includes factors that localize to cytoplasmic liquid-like condensates such as the P granule, the Z granule, or *Mutator* foci. These factors likely promote RNAi inheritance by acting with RdRPs to amplify siRNA populations each generation (Wan et al. 2018; Spracklin et al. 2017). The secondary category of factors are members of a nucleus-specific branch of the RNAi pathway (nuclear RNAi or NRDE pathway) (Buckley et al. 2012b; Ashe et al. 2012; Shirayama et al. 2012; Spracklin et al. 2017; Wan et al. 2018). According to current models of nuclear RNAi, AGOs bind and escort siRNAs to nuclei where these ribonucleoprotein complexes locate RNA Polymerase II (RNAP II)-dependent nascent transcripts based on complementarity to trigger co-transcriptional gene silencing (termed nuclear RNAi) (Guang et al. 2008a, 2010; Buckley et al. 2012b). HRDE-1 and NRDE-3 are two tissue-specific nuclear AGOs that drive nuclear RNAi in germ cells and somatic cells, respectively (Guang et al. 2008b; Buckley et al. 2012b; Ashe et al. 2012; Shirayama et al. 2012). The nuclear AGOs recruit downstream nuclear RNAi effectors (NRDE-1/2/4) to genomic sites of RNAi to direct histone post-translational modifications (PTMs) (*e.g*. H3K9me3 and H3K27me3) as well as inhibition of RNAP II during the elongation step of transcription via an unknown mechanism,(Guang et al. 2008b, 2010; Burkhart et al. 2011; Buckley et al. 2012a; Mao et al. 2015). While it is not yet clear why nuclear RNAi is needed for RNAi inheritance, it is known that nuclear RNAi is needed for transgenerational siRNA expression, suggesting that the co-transcriptional gene silencing and cytoplasmic siRNA amplification systems may be connected in some way.

If RdRP enzymes can amplify small RNA populations each generation during RNAi inheritance, then why doesn’t RNAi inheritance last forever? There could be fundamental biochemical limitations that prevent RNAi inheritance from lasting forever, or *C. elegans* might possess systems that actively prevent long-term inheritance. Recently, loss-of-function mutations in *met-2*, which is one of two *C. elegans* H3K9 methyltransferases responsible for depositing the majority of H3K9me2 found in *C. elegans*, were shown to cause RNA-directed TEI to last longer than normal (Andersen and Horvitz 2007; Towbin et al. 2012; Checchi and Engebrecht 2011; Lev et al. 2017). The reasons why loss of MET-2 enhances RNAi inheritance are not known, but may involve global alterations in gene expression that indirectly promote RNAi inheritance (Lev et al. 2017). To further our understanding of why RNAi inheritance doesn’t last forever, we conducted a forward genetic screen to identify additional factors that limit the duration of RNAi inheritance. Our screen identified the gene <u>h</u>eritable <u>e</u>nhancer of <u>R</u>NA<u>i</u> 1 (*heri-1*). We find that HERI-1 inhibits nuclear RNAi-based co-transcriptional gene silencing and this ability is likely the reason why loss of HERI-1 promotes RNAi inheritance. Interestingly, HERI-1 is physically recruited to genes undergoing nuclear RNAi, suggesting that HERI-1 may be a direct and dedicated regulator of nuclear RNAi and, therefore, TEI. Surprisingly, the recruitment of this negative regulator of nuclear RNAi to genes is itself dependent upon the nuclear RNAi. To explain these results, we propose that nuclear RNAi has both pro and anti-silencing functions and that the generational perdurance of RNAi inheritance is set by the relative contributions of these two opposing outputs.

## Results

### A genetic screen identifies <u>h</u>eritable <u>e</u>nhancers of <u>R</u>NA<u>i</u> (*heri*) genes

We conducted a genetic screen to identify factors that limit the generational perdurance of RNAi inheritance in *C. elegans* (Fig. 1). *gfp* RNAi silences a *gfp::h2b* reporter transgene for 6-9 generations (Vastenhouw et al. 2006; Buckley et al. 2012a). *oma-1(zu405ts*) is a temperature-sensitive (ts) gain-of-function (*gf*) lethal allele of *oma-1* (*Lin* 2003)*. oma-1* RNAi silences *oma-1(gf)* and, therefore OMA-1(GF)-mediated lethality for 4-7 generations (R. M. Alcazar, Lin, and Fire 2008; Buckley et al. 2012a). We EMS mutagenized *gfp::h2b*; *oma-1(gf)* animals and, two generations after mutagenesis, exposed animals to *gfp* and *oma-1* dsRNAs. In parallel, we propagated non-mutagenized control *gfp::h2b*; *oma-1(gf)* animals similarly treated with *gfp* and *oma-1* dsRNAs. Animals were propagated in the absence of further RNAi until non-mutagenized control animals failed to inherit *oma-1* or *gfp* silencing (≅7 generations). Strains that exhibited silencing for an additional seven generations were isolated and kept for further study. From approximately ~90,000 mutagenized genomes, we isolated twenty strains fulfilling these criteria. We refer to the genes mutated in these strains as the <u>h</u>eritable <u>e</u>nhancer of <u>R</u>NA<u>i</u> (*heri*) genes.

**Figure 1.**
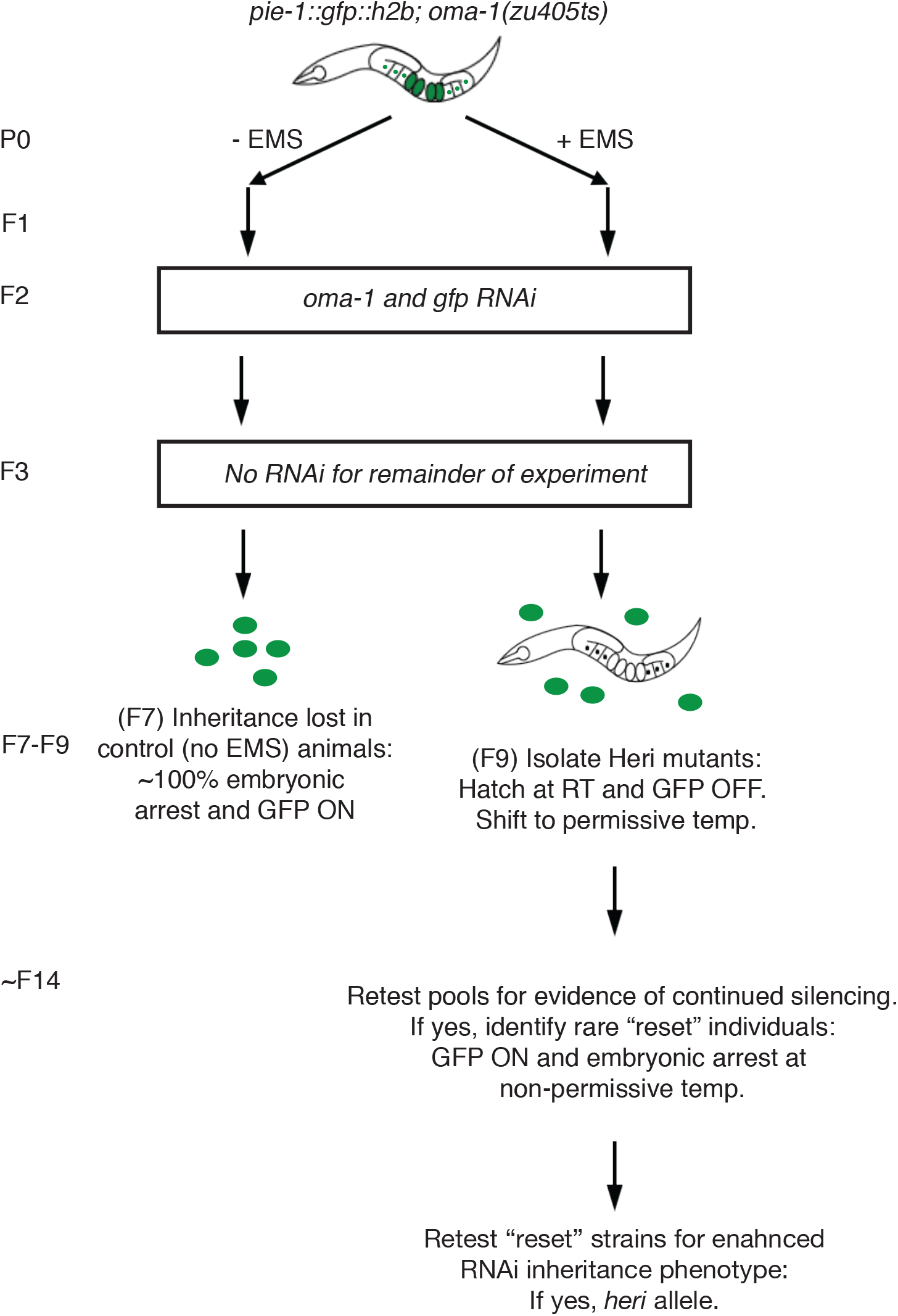
A genetic screen to identify <u>h</u>eritable <u>e</u>nhancers of <u>R</u>NA<u>i</u> (*heri*) genes. *pie-1::h2b::gfp* and *oma-1(zu405ts*) can be silenced heritably by RNAi (Vastenhouw et al. 2006; Alcazar et al. 2008). We treated *pie-1::h2b::gfp;oma-1(zu405ts*) animals with the mutagen ethyl methanesulfate (EMS). Control animals were left untreated. After two generations, animals were fed bacteria expressing double stranded RNA targeting *gfp* and *oma-1* simultaneously. Embryos from these animals were isolated and shifted to a non-RNAi (OP50) food source and grown at 20°C, which is the non-permissive temperature for *oma-1 (zu405ts*). Embryos were isolated for an additional 5 generations until control animals stopped inheriting *gfp* and *oma-1(zu405ts*) silencing (*gfp* ON and embryonic arrest, ~F7). Mutagenized pools were allowed to grow for 2 more generations and potential Heri strains (animals were *gfp* OFF and alive-indicating *pie-1::h2b::gfp* and *oma-1(zu405ts*) were still being silenced) were singled. After 5 additional generations, those populations still inheriting gene silencing were kept for further study. *heri* genes were identified by a combination of whole genome sequencing and bioinformatic analysis (see methods and Fig. S1). Rare animals within these populations had lost gene silencing at *pie-1::h2b::gfp* and *oma-1(zu405ts*). These animals were isolated to establish mutant lines that expressed reporter genes (termed “reset”).

### *heri-1* encodes a chromodomain protein with homology to Ser/Thr protein kinases

To assign identity to the *heri* genes, we used whole genome sequencing coupled with custom scripts and the bioinformatic software CloudMap (Minevich et al. 2012; Spracklin et al. 2017) (Fig. S1). This approach identified four independent mutations within the *C. elegans* gene *cec-9/c29h12.5* (Fig. 2A). For reasons outlined below we refer to *cec-9/c29h12.5* as *heri-1*. From strains harboring mutations in *heri-1*, we isolated rare individuals in which *gfp* and *oma-1* silencing had been lost and we used these “reset” strains to ask if these lines were indeed enhanced for RNAi inheritance. This analysis showed that RNAi inheritance after *oma-1* or *gfp* RNAi was enhanced in all four *heri-1* mutant strains (Fig. 2B and 2C). To confirm the molecular identity of *heri-1*, we tested an independently isolated allele of *heri-1* (*gk961392*) for RNAi inheritance. Note, *gk961392* likely represents a loss-of-function allele of *heri-1*, as *gk961392* is a deletion that removes conserved domains of HERI-1 (see below) and alters the reading frame (Fig. 2A). *gk961392* animals exhibited an enhanced RNAi inheritance phenotype (Fig. 2D and Fig. S2). We conclude that *cec-9/c29h12.5* is *heri-1* and that one function of HERI-1 is to limit the generational perdurance of RNAi inheritance.

**Figure 2.**
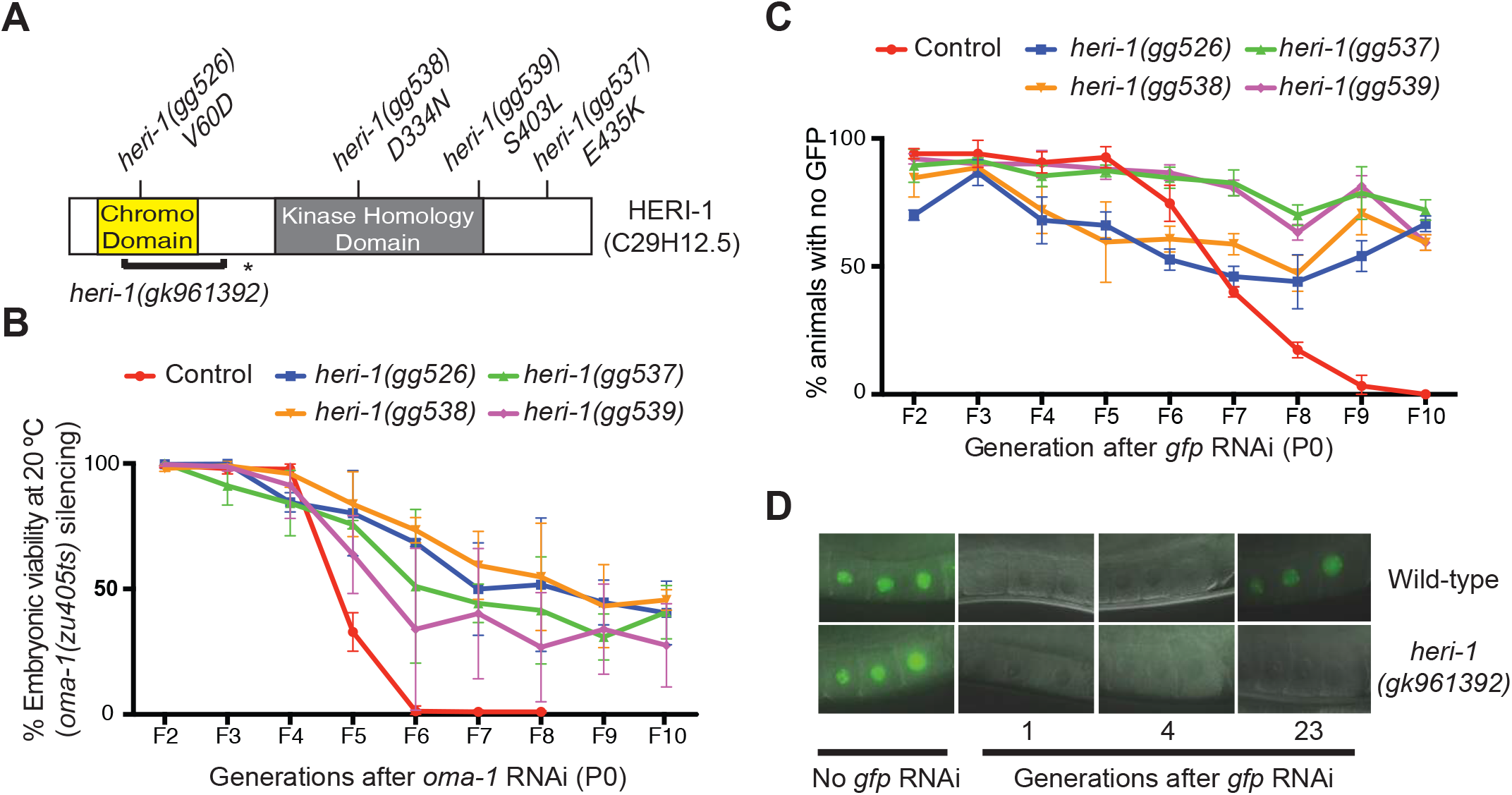
HERI-1 limits transgenerational epigenetic inheritance. **A)** Diagram of HERI-1 domain structure. *heri-1* alleles identified in our genetic screen and the *heri-1(gk961392*) deletion allele used in this study are indicated. Asterisk represents premature stop codon. **B)** RNAi inheritance assay (see methods) showing percent embryonic viability (*oma-1(zu405ts*) silencing) over generations after *oma-1* RNAi feeding in our starting strain (control, YY565) and *heri-1* mutants that were identified in the Heri screen and that had been “reset” for reporter gene expression (see Fig. 1). Error bars represent standard deviations of the mean. **C)** *gfp* inheritance assay showing the percentage of animals of the indicated genotypes showing *gfp* silencing in control and reset *heri-1* strains over generations after *gfp* RNAi exposure. **D)** Representative images of oocytes in wild type and *heri-1(gk961392*) animals that harbor the *pie-1::h2b::gfp* transgene during *gfp* RNAi inheritance assay. Generations after *gfp* RNAi are indicated.

### HERI-1 is expressed in germ cell nuclei

*heri-1* encodes a protein with two conserved domains, a chromodomain and a serine/threonine kinase-like domain (Fig. 2A). To understand more about HERI-1, we used CRISPR/Cas9 to introduce a C-terminal *gfp::3xflag* epitope to the 3’ terminus of the *heri-1* locus (Dickinson et al. 2015; Farboud and Meyer 2015; Paix et al. 2014; Arribere et al. 2014). In *heri-1::gfp::3xflag* animals, we observed HERI-1::GFP expression in the nuclei of adult germ cells (Fig. 3A). No HERI-1::GFP signal was observed in somatic tissues in adult animals (data not shown). GFP fluorescence was expressed diffusely throughout nuclei and did not colocalize with mitotic chromosomes (Fig. 3A and Fig. S3). Similar results were seen when CRISPR was used to introduce a *3xflag* epitope to *heri-1* and anti-FLAG immunofluorescence (IF) was used to detect HERI-1::FLAG (Fig. S3). RNAi inheritance assays indicated that CRISPR modified *heri-1* locus produced functional HERI-1 protein (Fig. S4). During embryogenesis in *C. elegans*, four asymmetric cell divisions partition germline determinants into the germline blastomeres termed the P_0_-P_4_ cells. During these early stages of embryogenesis, HERI-1::GFP was expressed in both germline and somatic blastomeres (Fig. S5 and data not shown). In the P1 blastomere, we detected a low level of GFP fluorescence in cytoplasmic puncta whose distribution and morphology were reminiscent of P granules (Fig. S5). By larval stage one (L1), HERI-1::GFP:: 3xFLAG was seen exclusively in the primordial germ cells Z2 and Z3, and was no longer seen in somatic tissues (Fig. 3B). HERI::GFP was expressed in germ cell nuclei throughout the remainder of germline development and no GFP expression could be seen in the soma during larval stages of development (Fig. 3B and data not shown). We conclude that HERI-1 is a germline-expressed protein that localizes predominantly to nuclei.

**Figure 3.**
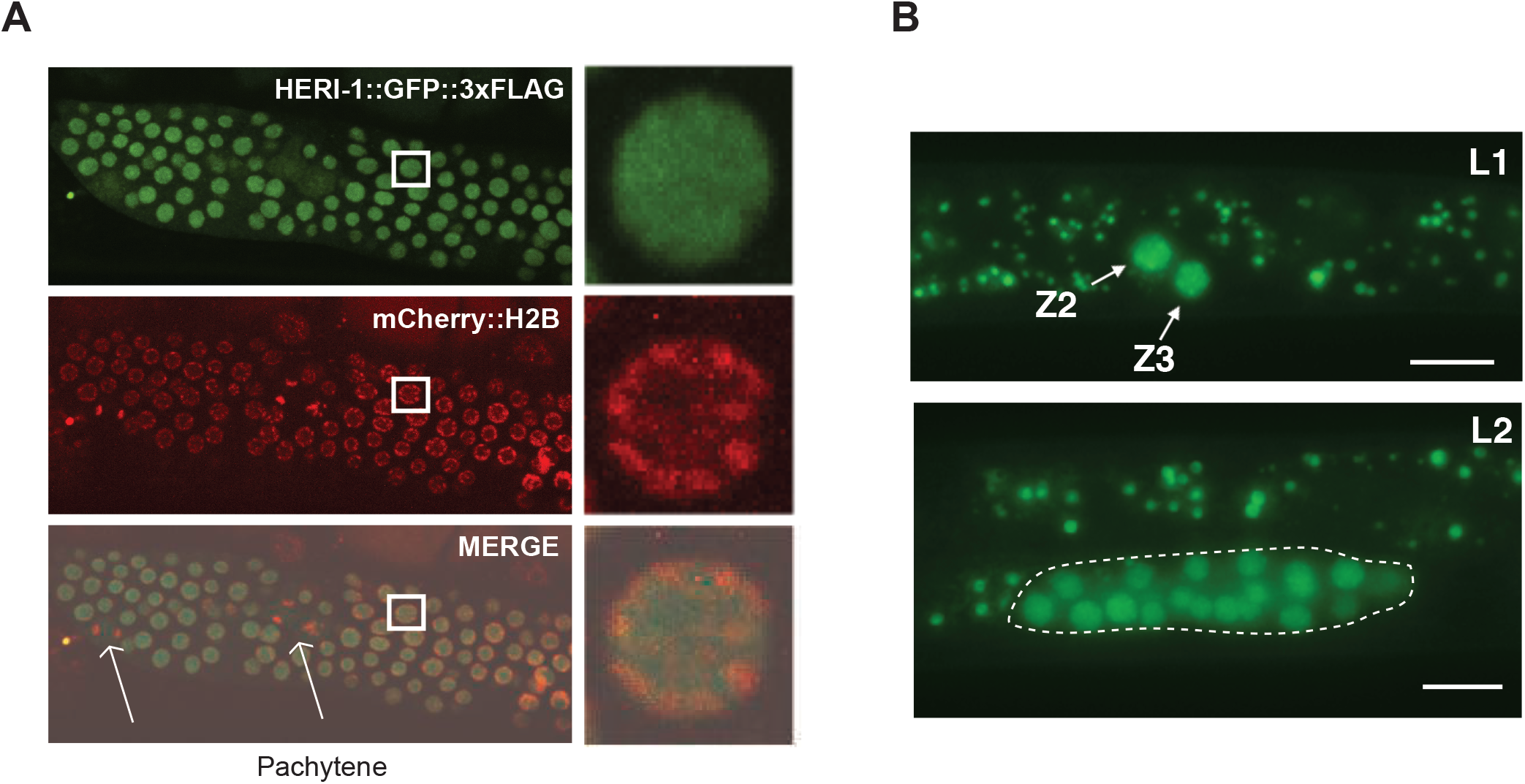
HERI-1 is expressed in germ cell nuclei. **A)** Micrographs of adult pachytene germline cells from animals expressing HERI-1::GFP::3XFLAG and mCHERY::H2B. Note HERI-1::GFP::3XFLAG does not appear to colocalize with chromatin in dividing cells (arrows). **B)** Micrograph of a larval stage one (L1) and two (L2) animals. HERI-1::GFP::3XFLAG is expressed in Z2 and Z3 in L1 and in the developing germline of L2 animals. In these animals, no HERI-1::GFP signal was observed in the soma. Note: non-germline fluorescence signal is due to autofluorescence, which is also present in wild-type animals. HERI-1::GFP is expressed throughout the remainder of germline development and no GFP fluorescence could be observed in somatic cells during this time (not shown).

### HERI-1 limits transgenerational siRNA and H3K9me3 expression

RNAi inheritance in *C. elegans* is correlated with both the heritable expression of siRNAs, which are anti-sense to genes undergoing heritable silencing, as well as with the heritable deposition of repressive histone marks (*e.g*. H3K9me3) on genes that are undergoing heritable silencing (Burton, Burkhart, and Kennedy 2011; Buckley et al. 2012a; Gu et al. 2012b; Ashe et al. 2012; Shirayama et al. 2012). Therefore, inherited siRNAs and/or H3K9me3 may be the informational vectors that drive RNAi inheritance. We asked if the enhanced RNAi we see in *heri-1* mutant animals was associated with an increase in the number of generations during which siRNAs and repressive chromatin marks were inherited. To do this, we used H3K9me3 ChIP and *gfp* siRNA Taqman probes to measure H3K9me3 deposited on *gfp* gene and *gfp* siRNA levels, respectively, in wild-type and *heri-1* mutant animals twenty generations after *gfp* RNAi had been initiated. Both H3K9me3 and siRNAs remained elevated in the F21 progeny of *heri-1* mutant animals exposed to dsRNA (Fig. S6), indicating that the enhanced RNAi inheritance phenotype shown by *heri-1* mutant animals is likely due to an enhancement of the molecular pathways mediating RNAi inheritance in wild-type animals.

### HERI-1 inhibits nuclear RNAi

What pathway(s) might HERI-1 regulate to limit RNAi inheritance? The *C. elegans* nuclear RNAi pathway is required for RNAi inheritance: in animals lacking components of the nuclear RNAi machinery, H3K9me3 is not deposited on chromatin, siRNAs are not heritably expressed, and gene silencing is not maintained in inheriting generations (Burton, Burkhart, and Kennedy 2011; Buckley et al. 2012a; Gu et al. 2012b; Ashe et al. 2012; Shirayama et al. 2012). The reason why nuclear RNAi is needed for RNAi inheritance is not yet known (see discussion). Given that nuclear RNAi mediates RNAi inheritance and HERI-1 is a nuclear protein, we wondered if HERI-1 might limit RNAi inheritance by inhibiting nuclear RNAi. The following lines of evidence indicate that this is the case. First, the nuclear RNAi machinery directs H3K9me3 deposition at genomic loci exhibiting sequence homology to RNAi triggers (Guang et al. 2010; Gu et al. 2012b). Using H3K9me3 ChIP to quantify H3K9me3 levels before and after RNAi, we found that RNAi triggered the deposition of ≅6x more H3K9me3 in *heri-1* mutant animals than what is seen in wild-type animals after RNAi (Fig. 4A). Second, exposure of *C. elegans* to dsRNA results in the nuclear RNAi-based silencing of unspliced nascent RNAs (pre-mRNA) exhibiting sequence homology to trigger dsRNAs (Guang et al. 2008a, 2010; Burton, Burkhart, and Kennedy 2011). Using qRT-PCR to quantify pre-mRNA levels before and after RNAi, we found that the degree to which RNAi silenced pre-mRNAs was greater in *heri-1* mutant animals than in wild-type animals (Fig. 4B). HRDE-1 is a germline Argonaute required for RNAi inheritance in wild-type animals. *hrde-1* was epistatic to *heri-1* for RNAi inheritance, suggesting that RNAi inheritance in *heri-1* mutant animals depends upon nuclear RNAi (Fig. 4C). We conclude that HERI-1 is a negative regulator of nuclear RNAi. We speculate that the inhibition of nuclear RNAi is the mechanism by which HERI-1 negatively regulates RNAi inheritance.

**Figure 4.**
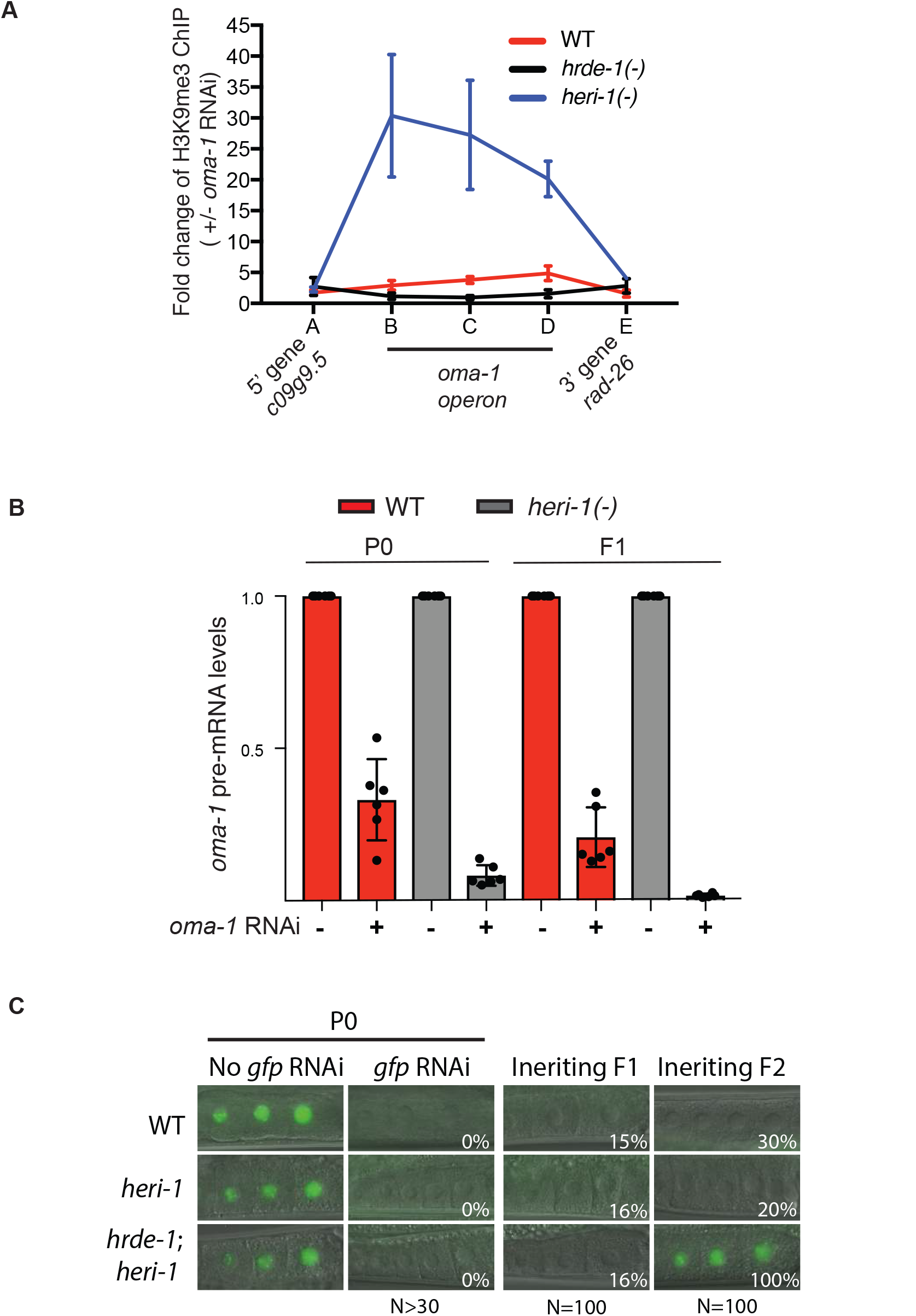
HERI-1 inhibits nuclear RNAi. **A)** H3K9me3 ChIP was conducted on wild type, *heri-1 (gk961392*), and *hrde-1(tm1200*) animals i n the progeny of animals exposed +/-to *oma-1* RNAi. Relative location of primers used for qPCR are indicated. Position along X-axis is not to scale with actual genomic loci (see Fig. 5). Data is from 3 biological replicates and error bars are standard deviations of the mean. Note, a value of one in this assay means RNAi had no effect on state of H3K9me3 at this locus. Data show that H3K9me3 is enriched within the *oma-1* operon after *oma-1* RNAi. *oma-1* operon contains three genes; *oma-1*, *spr-2*, and *c27b76.2*. Primer set B and C are in *oma-1*. Primer set D is in *spr-2*. **B)** Quantitative PCR (qPCR) data showing that the *oma-1* pre-mRNA is more silenced in *heri-1(gk961392*) animals than in wild-type animals after *oma-1* RNAi. Error bars are standard deviations of the mean. Note, a value of one would indicate that RNAi had no effect on *oma-1* pre-mRNA levels for this experiment. **C)** Animals of the indicated genotypes (*heri-1(gk961392*), and *hrde-1(tm1200)*) and expressing *pie-1::gfp::h2b* were exposed to +/-*gfp* RNAi. Representative micrographs of -1,-2, and -3 oocytes are shown. Percentages of animals inheriting silencing in the indicated generations are shown.

### RNAi directs HERI-1 to chromatin

Chromodomains can interact with post translationally modified (PTM) histones such as H3K9me3 and H3K27me3 (Eissenberg 2012). Nuclear RNAi in *C. elegans* directs H3K9me3 and H3K9me3 (Guang et al. 2010; Gu et al. 2012a; Buckley et al. 2012a; Mao et al. 2015). These observations hint at the possibility that HERI-1 might be physically recruited to genes undergoing nuclear RNAi and/or RNAi inheritance. To test this idea, we used HERI-1 chromatin IP (ChIP) to ask if RNAi would cause HERI-1 to associate with the chromatin of genes we had targeted by RNAi. Indeed, we found that, after *oma-1* RNAi, HERI-1::GFP::FLAG associated with the *oma-1* locus, but not the genes flanking *oma-1* (Fig. 5). Similar results were seen when animals expressing HERI-1::FLAG were treated with *oma-1* RNAi (Fig. S7). We conclude that RNAi can direct HERI-1 to interact with chromatin of genes that exhibit sequence homology to dsRNA triggers. The physical recruitment of HERI-1 to genes targeted by RNAi suggests that HERI-1 may play a direct role in inhibiting nuclear RNAi and, therefore, RNAi inheritance.

**Figure 5.**
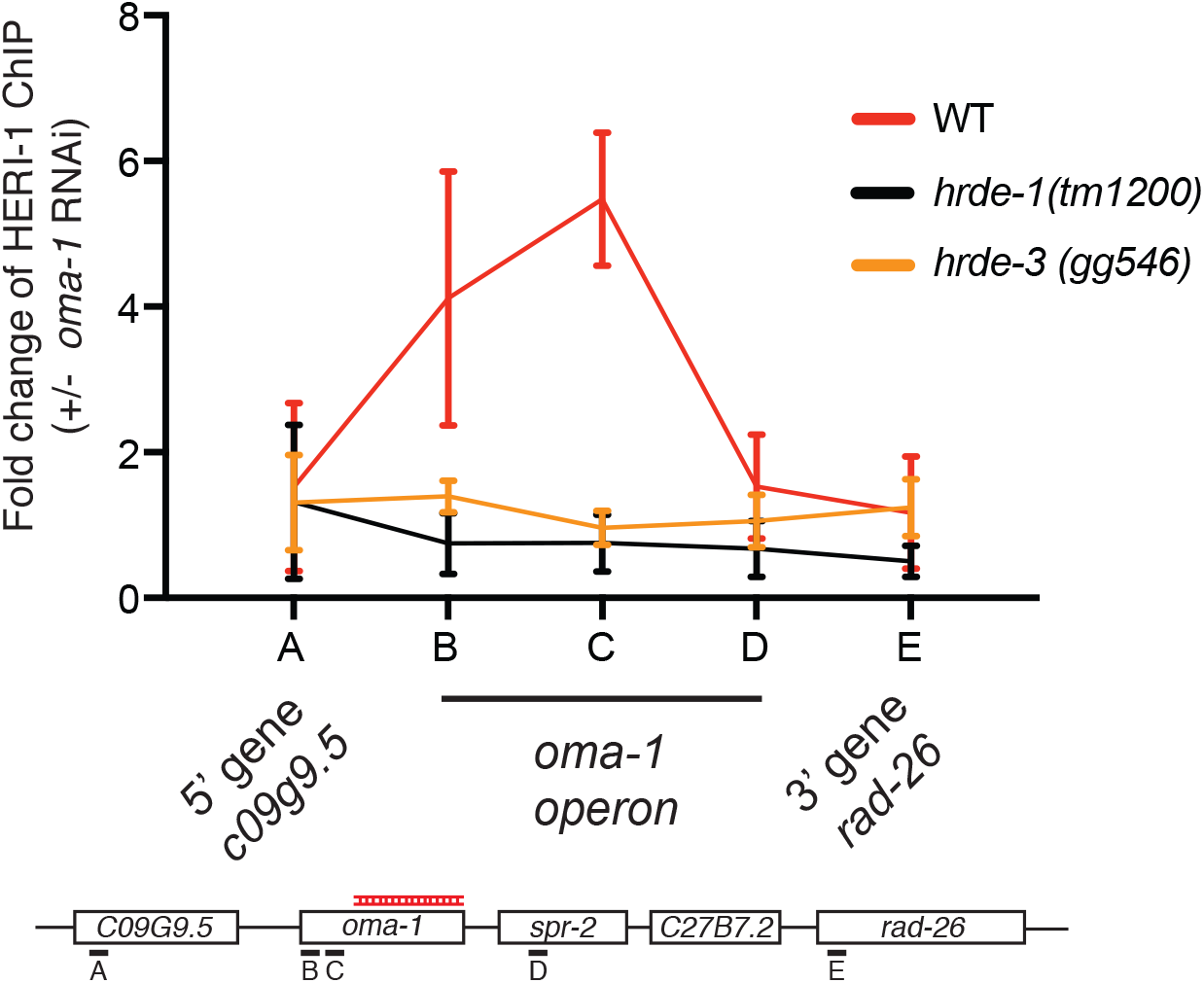
RNAi causes HERI-1 to associate with chromatin of genes undergoing heritable silencing. HERI-1::GFP::3xFlag animals were exposed to +/-*oma-1* RNAi and HERI-1 ChIP was conducted on progeny. Fold enrichment of HERI-1 on the *oma-1* locus, as well as neighboring loci, is shown and is expressed as a ratio of HERI-1 ChIP signals +/-*oma-1* RNAi. Data is from 3 biological replicates and error bars are standard deviations of the mean. Similar results were obtained when HERI-1::3xFLAG animals were subjected to a similar analysis (Fig. S7). A value of one would indicate that RNAi had no effect on HERI-1 ChIP. Note: primer set D does not show RNAi-induced HERI-1 binding, but did show an increase in H3K9me3 in Fig. 4 (see discussion).

### Recruitment of HERI-1 to chromatin requires nuclear RNAi

How is HERI-1 recruited to genes targeted by RNAi? *heri-1* encodes a chromodomain (Fig. 2A). Chromodomain proteins are reported to interact with H3K9me3 and H3K27me3 (Eissenberg 2012). Additionally, our genetic screen identified a missense mutation in the *heri-1* chromodomain (V60D), suggesting that this domain is important for HERI-1 function (Fig. 1A). For all these reasons, we wondered if nuclear RNAi-directed chromatin modifications (such as H3K9me3 or H3K27me3) might be responsible for recruiting HERI-1 to genes after RNAi. Note: such a scenario would be surprising, as nuclear RNAi normally promotes gene silencing while HERI-1 limits this silencing. HRDE-1 is a nuclear AGO that drives nuclear RNAi in germ cells (Buckley et al. 2012a). We exposed wild-type or *hrde-1(-)* animals to *oma-1* RNAi and used HERI-1 ChIP to quantify HERI-1 chromatin interactions in these animals. The analysis showed that HRDE-1 was required for RNAi to direct HERI-1 to bind chromatin, suggesting that nuclear RNAi is, indeed, needed to recruit HERI-1 to chromatin (Fig. 5). The putative H3K9 methyltransferase SET-32/HRDE-3 is, at least in some cases, required for RNAi inheritance as well as for RNAi-directed H3K9 methylation at the *oma-1* locus following *oma-1* RNAi (Ashe et al. 2012; Spracklin et al. 2017). We found that SET-32/HRDE-3 was also required for *oma-1* RNAi to recruit HERI-1 to the *oma-1* gene, hinting that H3K9me3 may be a component of the chromatin signature that recruits HERI-1 to chromatin (Fig. 5). [Note: *heri-1* chromodomain mutations destabilize HERI-1, making additional obvious tests of this model difficult (see discussion).] We conclude that HRDE-1 and SET-32 are required for recruiting the negative regulator of nuclear RNAi HERI-1 to genomic sites of RNAi. Because HRDE-1 and SET-32 are themselves required for nuclear RNAi, the data suggest that the nuclear RNAi machinery in *C. elegans* has both pro- and anti-silencing outputs. First, the machinery links inherited siRNAs to co-transcriptional gene silencing. Second, the machinery places limits on this silencing by recruiting the negative regulator HERI-1 to sites of nuclear RNAi, perhaps by altering chromatin landscapes in ways recognized by the HERI-1 chromo-domain.

### *heri-1* mutant animals have defective sperm, which may be caused by hyperactive nuclear RNAi

While working with *heri-1* mutant animals, we noticed that ≅20% of *heri-1(gk961392)* animals had a single or both gonad arms containing more oocytes than normal (termed a stacked oocyte phenotype) (Fig. 6A/B). Animals harboring any one of our four other *heri-1* mutations also showed this stacked oocyte phenotype, albeit at a somewhat reduced rate from animals harboring the *heri-1* deletion allele (Fig. S8). These data indicate that the loss of HERI-1 causes oocytes to accumulate within *C. elegans* germlines. Stacked oocytes are often seen in *C. elegans* lacking functional sperm (Schedl and Kimble 1988). Two additional lines of evidence support the idea that *heri-1* mutant animals have dysfunctional sperm. First, *heri-1(-)* animals with stacked oocytes were sterile (Fig. 6C). Second, wild-type sperm (introduced by mating) rescued the fertility defects of *heri-1* animals that had stacked oocytes (Fig. 6C). We conclude that ≅20% of animals that lack HERI-1 produce dysfunctional sperm or have defects in spermatogenesis.

**Figure 6.**
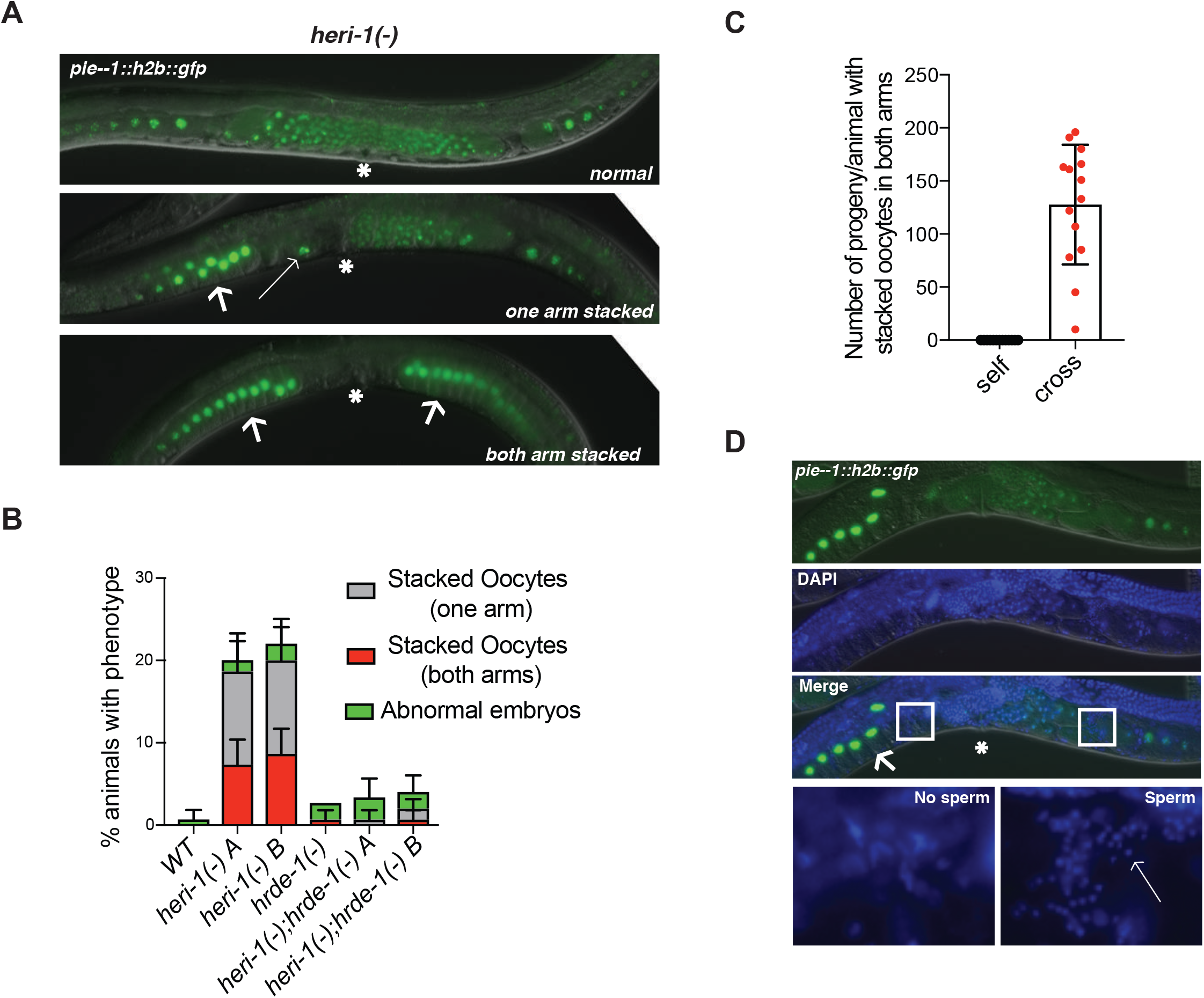
*heri-1* mutant animals have defective sperm. **A)** Fluorescent micrograph of wild-type or *heri-1(gk961392*) oocytes expressing *pie-1p::h2b::gfp*. 20% of *heri-1* mutant animals show a stacked oocytes in either one or both gonad arms. Asterisk indicates vulva. Thick arrows indicate stacked oocytes. Thin arrow indicate an unfertilized oocyte. **B)** Quantification of the indicated oocyte defects in *wild type, heri-1(gk961392), hrde-1(tm1200*), and double mutants animals. Error bars are standard deviations of the mean. **C)** Number of progeny from *heri-1(gk961392*) animals that had stacked oocytes in both gonad arms that were selfed (self) or after crossing to wild type (N2) males (cross). n=13 for self and n=14 for cross. **D)** DAPI staining of a *heri-1(gk961392);pie-1::h2b::gfp* animals that had stacked oocytes in one gonad arm. A magnification of spermatheca from both gonad arms (indicated by box in “merge” panel) is shown below. Arrow indicates gonad arm with stacked oocytes. Asterisk indicates vulva.

To try and better understand what goes wrong in *heri-1(-)* sperm, we isolated and DAPI stained animals in which one of two gonad arms contained stacked oocytes (Fig. 6D). Spermatheca from “normal” germline arms possesed sperm, however; in spermatheca from defective germline arms (with stacked oocytes), no sperm could be seen (Fig. 6D). The data suggest that *heri-1* sperm are defective because either: 1) they never develop, or 2) develop-but are dysfunctional and, consequently, are swept out of the spermatheca by oocytes transiting the spermatheca. Additional studies will be needed to differentiate between these two possibilities. We wondered if *heri-1(-)* sperm defects were related to HERI-1’s role in limiting nuclear RNAi. If so, a *hrde-1* mutation would be expected to suppress *heri-1(-)* sperm defects, as nuclear RNAi should no longer be hyperactive in animals that lack the AGO that directs nuclear RNAi. To test this idea, we quantified sperm defects in *heri-1*(-), *hrde-1*(-), and *heri-1*(-); *hrde-1(-)* double mutant animals and found that, indeed, *hrde-1* was epistatic to *heri-1* with regards to sperm defects (Fig. 6B). The data are consistent with the idea that HERI limits nuclear RNAi (directed by nuclear siRNAs and HRDE-1) in sperm nuclei and that this regulation is needed for normal sperm function and/or development.

## Discussion

*heri-1* encodes a chromo/kinase domain protein that negatively regulates nuclear RNAi and RNAi inheritance. We speculate that it is the ability of HERI-1 to limit nuclear RNAi that allows HERI-1 to limit RNAi inheritance. Interestingly, we find that HERI-1 is recruited to genes targeted by RNAi, suggesting that HERI-1 may be a direct regulator of RNAi inheritance, and, therefore, that *C. elegans* possess genetically encoded systems dedicated to limiting RNA-directed TEI during normal reproduction. Finally, the NRDE nuclear RNAi system, which is itself needed for nuclear RNAi, is needed for localizing the negative regulator of nuclear RNAi HERI-1 to genes. These data suggest that the nuclear RNAi machinery has two functions during RNAi inheritance: 1) to impart epigenetic marks that promote co-transcriptional gene silencing and 2) to recruit negative regulators of nuclear RNAi in order to limit the number of generations over which this epigenetic information is inherited.

### How does HERI-1 inhibit nuclear RNAi?

We do not understand how HERI-1 limits nuclear RNAi. HERI-1 has homology to serine/threonine protein kinases, and one of the *heri-1* alleles we identified in our genetic screen (*gg538*) alters a conserved aspartate residue within the HERI-1 kinase-like domain (Fig. 2A). Thus, the HERI-1 kinase-like domain is likely important for HERI-1 to inhibit nuclear RNAi and, therefore, RNAi inheritance. HERI-1 is unlikely to be an active protein kinase, however, as its kinase domain lacks active site residues required for kinase activity in related protein kinases (Nolen, Taylor, and Ghosh 2004). For instance, the GxGxxG, VAIK, and DFG motifs, which contribute to Mg^2+^ and ATP binding in canonical protein kinases, are not conserved in HERI-1 (Fig. S9). Additionally, we failed to detect kinase activity associated with recombinant HERI-1 protein *in vitro* (Fig. S10). Thus, we predict that HERI-1 is a pseudokinase. Pseudokinases are fairly common in eukaryotic genomes where they often play important roles in diverse aspects of cellular physiology, such as acting as scaffolds, anchors, or allosteric regulators of other proteins, including other protein kinases (Eyers and Murphy 2013). Given that HERI-1 is recruited to sites of nuclear RNAi, we speculate that HERI-1 negatively regulates nuclear RNAi by interacting with and regulating other pro-silencing factors at sites of nuclear RNAi (Fig. 7). Potential targets of such regulation include the nuclear RNAi factors HRDE-1 and NRDE-1/2/4 as well as the chromatin modifying enzymes that impart repressive modifications to chromatin in response to nuclear RNAi (Guang et al. 2010; Burkhart et al. 2011; Buckley et al. 2012a) (Fig. 7). HERI-1 IP-MS experiments might allow the regulatory targets of HERI-1 to be identified.

**Figure 7.**
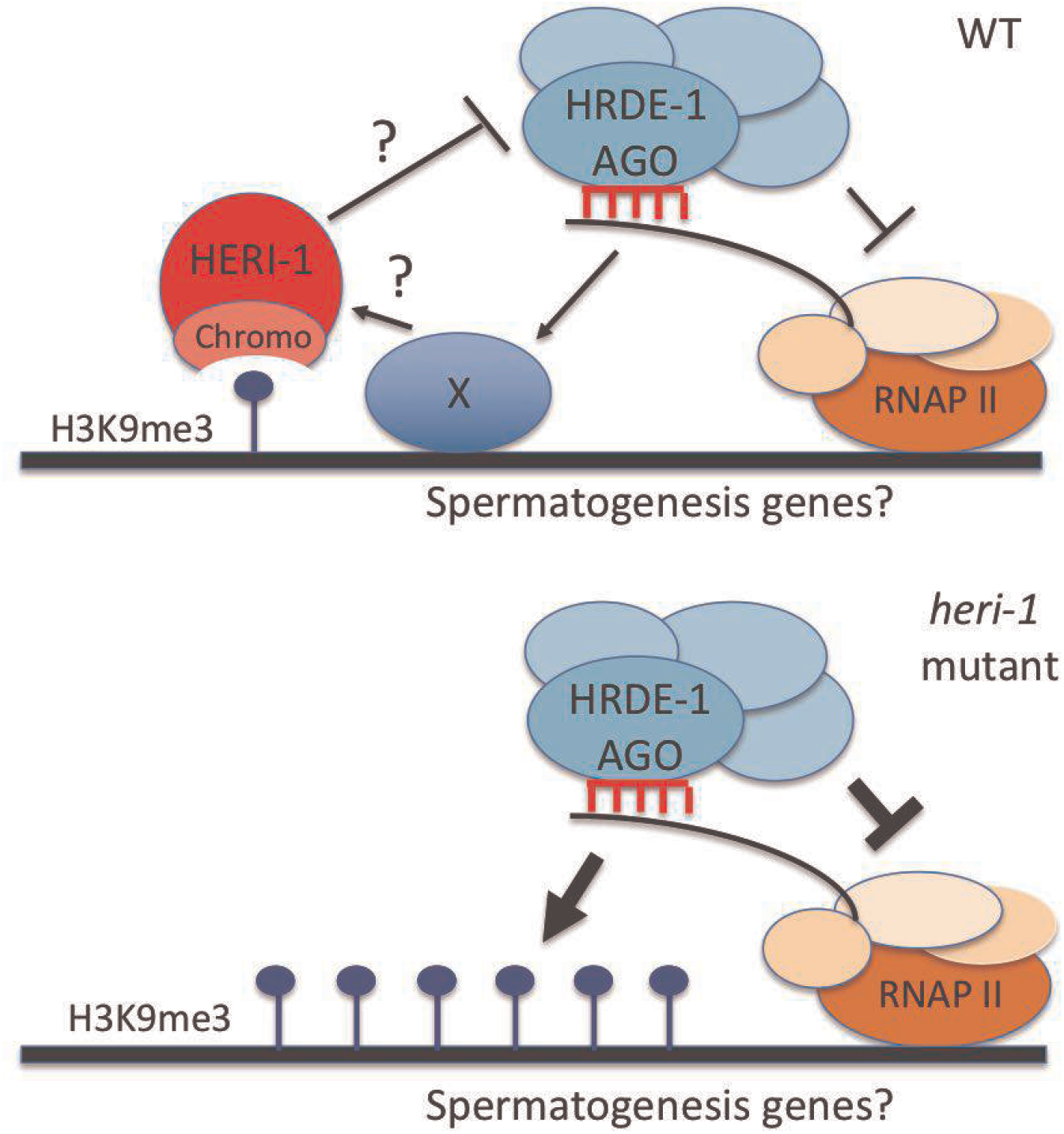
Model for the role of HERI-1 in limiting RNA inheritance. The AGO HRDE-1 uses siRNAs as guides to recognize and bind pre-mRNAs with sequence homology to siRNAs. HRDE-1 then recruits downstream silencing factors, such as the NRDEs (unlabeled teal ovals) and chromatin-modifying enzymes, to deposit H3K9me3 as well as another, currently unknown (X), signal on chromatin to mark genes undergoing nuclear RNAi. Note: X could be another chromatin PTM or a chromatin/DNA binding protein. X and H3K9me3 cooperate to recruit HERI-1 to chromatin. HERI-1 is a pseudokinase that may limit RNAi inheritance by binding and regulating pro-silencing proteins (such as HRDE-1 or NRDEs) that are present on genes undergoing nuclear RNAi/RNAi inheritance. The model predicts that nuclear RNAi drives co-transcriptional gene silencing by inhibiting RNA Polymerase II (RNAPII) while, at the same time, limiting this silencing by recruiting negative regulators of the process, such as HERI-1, to sites of nuclear RNAi/RNAi inheritance.

### How is HERI-1 recruited to chromatin?

HERI-1 is recruited to chromatin in response to RNAi (Fig. 5). Recruitment is dependent upon the nuclear RNAi AGO HRDE-1, suggesting that nuclear siRNAs direct HERI-1 recruitment to chromatin (Fig. 5). How might nuclear siRNAs recruit HERI-1 to chromatin? The nuclear RNAi pathway directs the deposition of repressive chromatin marks such as H3K9me3 and H3K27me3 on the chromatin of genes undergoing nuclear RNAi (Guang et al. 2010; Burkhart et al. 2011; Gu et al. 2012a; Mao et al. 2015). Additionally, HERI-1 has a chromodomain, and chromodomains are known to bind post-translationally modified (PTM) histones, such as H3K9me3 and H3K27me3 (Eissenberg 2012). Together, these observations suggest the following model: First, nuclear RNAi directs PTMs such as H3K9me3 on chromatin. Second, these PTMs interact with the chromodomain of HERI-1 to recruit HERI-1 to genes so that HERI-1 can regulate nuclear RNAi. Consistent with the model, we find that the putative H3K9 methyltransferase SET-32, which is required for RNAi-directed H3K9 methylation in germ cells, is required for RNAi to direct HERI-1 to chromatin (Spracklin et al. 2017) (Fig. 7).

Two observations indicate that H3K9me3 is unlikely to be sufficient to recruit HERI-1 to chromatin, however. First, *oma-1* is in an operon with *spr-2*. We detected an increase in H3K9me3, but not an increase of HERI-1, on the *spr-2* gene after *oma-1* RNAi (Fig. 4/5, primer D). Thus, H3K9me3 may be necessary for recruiting HERI-1 to chromatin, but it cannot be sufficient. We speculate that other, currently unknown, signals (*e.g*. additional chromatin PTMs), which are present on the *oma-1* gene, but not the *spr-5* gene after RNAi, may act with H3K9me3 to localize HERI-1 to *oma-1* chromatin. Second, animals lacking the putative H3K9me3 methyltransferase SET-32/HRDE-3 (in which HERI-1 fails to localize to chromatin) are defective, not enhanced, for RNAi inheritance (Ashe et al. 2012; Spracklin et al. 2017). Thus, the function of H3K9me3 in nuclear RNAi (and RNAi inheritance) cannot be limited to the recruitment of negative regulators of co-transcriptional gene silencing, such as HERI-1 to chromatin. How H3K9me3 promotes nuclear RNAi and RNAi inheritance in *C. elegans* remains surprisingly mysterious as studies addressing this question have generated somewhat contradictory results. For instance, genetic screens, and candidate gene approaches, have shown that H3K9me3-like enzymes (*e.g*. SET-32) are required (at least in some cases) for RNAi-directed H3K9 methylation and for RNAi inheritance (Ashe et al. 2012; Spracklin et al. 2017). On the other hand, *C. elegans* lacking two other H3K9 methyltransferases, MET-2 and SET-25, lack biochemically detectable levels of H3K9me2/3 and yet do not show obvious defects in RNAi inheritance (at least in some assays) (Towbin et al. 2012; Kalinava et al. 2017). Indeed, depletion of one of these enzymes (MET-2) actually enhances RNAi inheritance (at least in some assays) (Lev et al. 2017). We speculate some of this confusion may relate to the fact that H3K9me2/3 may have opposing functions during RNAi inheritance; 1) the promotion of gene silencing via heterochromatin-like systems and 2) the inhibition of gene silencing via recruitment of negative regulators like HERI-1. If the relative contributions of these two opposing pathways differed between genes and across development, this could help explain why understanding the role of H3K9me3 in RNAi inheritance has been so difficult. Biochemical studies to ask if the HERI-1 chromodomain actually binds H3K9me3 (or some other PTM) will be needed to definitively test the model. Analysis of additional *heri* genes, identified by our genetic screen, could also test the model by identifying factors and pathways that direct HERI-1 to chromatin in response to RNAi. Note, these studies will need to account for fact that mutations in the HERI-1 chromodomain destabilize HERI-1 protein, suggesting that HERI-1/chromatin interactions are likely important for HERI-1 stability (Fig. S11).

### HERI-1 and sperm function

Why would *C. elegans* need factors that limit nuclear RNAi in germ cells? *C. elegans* expresses an abundant class of endogenous (endo) siRNAs that are thought to direct nuclear RNAi in nuclei during the normal course of growth and reproduction (Lee, Hammell, and Ambros 2006; Guang et al. 2008b, 2010; Burkhart et al. 2011; Billi, Fischer, and Kim 2014). Some of these endo siRNAs are thought to be present in sperm progenitors where they regulate gene expression programs needed for sperm development and function (Sijen et al. 2001; Kennedy, Wang, and Ruvkun 2004; Duchaine et al. 2006; Pavelec et al. 2009; Gent et al. 2009, 2010; Conine et al. 2010; Vasale et al. 2010). We find that *heri-1* mutant animals exhibit sperm defects, which are suppressed by loss of the germline nuclear RNAi AGO HRDE-1. The data suggest that one function of TEI-regulating systems in *C. elegans* is to limit endogenous nuclear RNAi (directed by sperm 22G siRNAs) in mature sperm, or sperm progenitors, and that this gene regulation is important for sperm function. Such gene misregulation might occur at genes normally targeted by nuclear 22G endo siRNAs in wild-type animals or, alternatively, at genes that are not normally subjected to nuclear RNAi. Regarding this latter model, endogenous RNAi (like experimental RNAi) is driven by 22G siRNAs, which are maintained over generations by RdRP enzymes (Gent et al. 2009, 2010; Buckley et al. 2012a). Such a self-amplifying feed-forward mode of gene regulation could be dangerous if genes that are normally not silenced by endo siRNAs or are only marginally silenced to inappropriately enter this heritable silencing pathway. We speculate that HERI-1 may promote germ cell health by preventing and/or reversing such unwanted and runaway heritable gene silencing. The identification of HERI-1 regulated sperm genes (via HERI-1 ChIP), and/or loci regulated by nuclear RNAi in wild-type and *heri-1* mutant animals (H3K9me3 ChIP or RNA-Seq), should allow this idea to be tested.

## Acknowledgements

We would like to thank the members of the Kennedy lab for helpful discussions of the data and the manuscript.

## No conflict of interest

The authors declare no conflict of interest.

## Materials and Methods

### Strains

For a complete list of strains used in this study, see supplemental table 1. Some strains were provided by the CGC, which is funded by NIH Office of Research Infrastructure Programs (P40 OD010440). All strains were maintained using standard laboratory conditions.

### *heri* screen

Larval stage 4 (L4) worms carrying both the *oma-1(zu405*) allele and the *pie-1::gfp::h2b* transgene were mutagenized with ethyl methanesulfate (EMS) at a concentration of 47mM for 4hrs at room temperature. Control worms were not exposed to EMS. Mutagenized and control worms were washed 3x with M9 buffer, and allowed to recover for 2hrs at 20°C in NGM plates with OP50. F2 progeny of mutagenized and control worms were isolated by hypochlorite treatment and exposed to *oma-1* and *gfp* RNAi simultaneously in 10cm RNAi plates at 20°C. In total, ~90,000 mutagenized genomes were screened in 74 independent pools. In the following generation embryos were isolated with hypochlorite treatment and fed OP50 at 20°C. We repeated this process until the control worms failed to inherit *oma-1* and *gfp* silencing (F7). In the F9, we isolated mutants that continued to silence *gfp* and *oma-1*. Five generations later, lines that continued to silence reporter genes were kept for further study. To identify candidate *heri* genes, we used whole genome sequencing coupled with the bioinformatics tool CloudMap as described in (Minevich et al. 2012). In short, genomic DNA was purified from control and mutants strains. Next generation sequencing libraries were prepared and sequenced using the Illumina HiSeq platform. Each genome was sequenced using paired-end sequencing at a coverage of about 20X. CloudMap software was used to identify, map and compare SNP’s between *heri* mutant strains. Genes that contained three (or greater) unique SNPs were kept as potential *heri* genes.

### RNAi Inheritance assays

For *gfp* inheritance assays: embryos were collected by hypochlorite treatment and placed in either control (HTT115) or *gfp* RNAi expressing bacteria at 20°C. The F1 embryos were collected again by hypochlorite treatment and fed OP50. When animals reached gravid adult stage, they were scored for germline *gfp* expression in groups of 50 worms per biological replicate. For Figure 2C and D, this process was repeated for 10 and 23 generations, respectively. For *oma-1* inheritance assays in Figure 1B: Embryos were collected by hypochlorite treatment and placed in either control (HTT115) or *oma-1* RNAi expressing bacteria at 20°C. The F1 embryos were collected again by hypochlorite treatment and fed OP50. For each biological replicate and for each generation, we singled 6 individual at the L4 stage of development and 3 days later we counted the number of hatched embryos as well as arrested embryos. The percent viability (*oma-1* RNAi inheritance) of each strain was calculated as the number of embryos that hatched within 24 hours over the total number of laid eggs.

### Construction of HERI-1 tagged strains using CRISPR

For construction of the C-terminal *heri-1::3xflag* strain: we used the Co-CRISPR strategy described in (Arribere et al. 2014). In short, candidate guide RNAs were calculated using the CRIPSR design tool at Massachusetts Institute of Technology (MIT) (crispr.mit.edu). We chose the guide RNA sequence 5’-TCATGCGAAACGAGAGAAAG-3’ because it’s PAM sequence of GTGG is located 4 nucleotides upstream of *heri-1’s* termination codon, and it had low off-target effect predictions. As a repair template, we used a single stranded DNA oligo (4nM ultramer HPLC purified from IDT) which contained 50bp of homology on each side of *heri-1’s* termination codon, and it had the PAM sequence mutated to GTTG to prevent repeated Cas9 cutting. Germlines of gravid hermaphrodites were injected with the following injection mix: gRNA (20ng/μl), repair template (20ng/μl), Co-CRIPSR markers pDD162(50ng/μl)(Addgene #47549), *unc-58* gRNA(20ng/μl), and AF-JA-76 (20ng/μl) all in 1X taq buffer (NEB). Uncoordinated (Unc) progeny of injected worms were singled, allowed to self-fertilized, lay progeny, and then were genotyped for the presence of the 3xflag insertion at the 3’end of *heri-1*. For the construction of the C-terminal *heri-1::gfp::3xflag* strain and the *heri-1::chromomutant:: gfp::3xflag*, we followed the CRISPR/Cas9 homologous recombination protocol described in (Dickinson et al. 2015). Details of this approach can be found here (http://wormcas9hr.weebly.com/). Repair template had ~500bp of homology to *heri-1* and contained a self excising cassette (SEC) with 3 features: a Hygromycin resistance gene, a Roller (Rol) marker gene and an heat shock inducible Cre recombinase gene. The SEC is flanked by LoxP sites. Gravid hermaphrodites were injected with an injection mix containing the following: 50ng/uL of pDD162 plasmid containing the Cas9 and sgRNA (Addgene #47549), 10ng/uL of repair template, 10 ng/μL pGH8 (*Prab-3::mCherry* neuronal co-injection marker; Addgene #19359), 5 ng/μL pCFJ104 (*Pmyo-3::mCherry* body wall muscle co-injection marker; Addgene #19328), 2.5 ng/μL pCFJ90 (*Pmyo-2::mCherry* pharyngeal co-injection marker; Addgene #19327). Injected worms (3 per plate) were incubated at 25°C for 2-3 days. Then, Hygromycin was added directly to the plates at a concentration of 250ug/mL. Plates were incubated at 25°C for 4 days. Animals that were Hygro resistant, Rol, and lacked the red fluorescent markers were isolated. To excise SEC, L1 animals were heat-shocked at 34°C for 4-5 hrs and shifted to 20°C for 3-5 days. Non-Rol animals were isolated and genotyped by PCR and Sanger sequencing for the *gfp::3xflag* insertion.

### Chromatin Immunoprecipitation and qPCR

We isolated embryos by hypochlorite treatment and fed larval animals HTT115 bacteria expressing *oma-1* dsRNA. The following generation, embryos were again collected by hypochlorite treatment and were fed OP50 (non-RNAi bacteria). Gravid hermaphrodites were collected and ~100uL pellet of packed worms were washed 2x with 1mL of M9 buffer, and then flashed frozen on liquid nitrogen in a 1.5mL tube. The pellet was crosslinked by incubating in 1mL of M9 buffer with 2% formaldehyde and allow to rotate for 30mins. Crosslinking reaction was quenched by addition of 54uL of 2.5M glycine and samples were then rotated for an additional 5mins. Animals were then washed 2x with 3mL of M9 buffer andresuspended in 0.5mL of FA lysis buffer (50mM Tris/HCl pH 7.5, 1mM EDTA, 1% Triton X-100, 0.1% sodium deoxycholate, 150mM NaCl) supplemented with complete Mini protease inhibitors (Roche). Animals were sonicated in a Qsonica Q800R sonicator, for 25 mins with a cycle of 30sec On, 30sec Off and 70% output. Insoluble debri was cleared by a quick spin in a microcentrifuge of 2mins at 5000rpm at 4°C. Extracts were transferred to new tubes, centrifuged at 15,000 rpm for 10min at 4°C. Protein concentration were calculated with Bradford assay (Biorad). Equal amounts of protein extracts were pre-cleared using 15ul of 50:50 slurry of unblocked protein A or G agarose beads (Millipore) in FA lysis buffer for 15mins at 4°C, while rotating. Extract/bead mixtures were centrifuged for 30sec at 3000rpm and supernatants were transferred to new tubes. For H3K9me3 ChIP, we used 2ug of Anti-H3K9me3 antibody (07-523 Millipore) and for HERI-1::3xFLAG ChIP we used 3ug of M2 anti-flag antibody (F1804 Sigma). Antibodies were added and incubated overnight at 4°C, while rotating. The following day, 20uL of protein A (for H3K9me3) and protein G (for Flag) were added to the extracts and allowed to bind for 2hrs at 4°C, while rotating. Beads were washed 2x with FA lysis buffer, 2x with FA lysis+500mM NaCl (50mM Tris/HCl pH 7.5, 1mM EDTA, 1% Triton X-100, 0.1% sodium deoxycholate, 500mM NaCl), 1x with LiCl buffer (0.25M LiCl, 1% NP-40, 1% sodium deoxycholate, 1mM EDTA, 10mM Tris/HCl pH 8.0) and 2x with TE buffer (1mM EDTA, 10mM Tris/HCl pH 8.0). All washes were performed in 1mL volumes and 1 min rotation at room temperature, beads were concentrated by centrifugation for 30sec at 3000rpms. Immunoprecipitated chromatin was eluted by incubatingbeads/ChIPs in 500uL of 0.1M Sodium Carbonate and 1% SDS for 30mins, while rotating at room temperature. Beads were concentrated by centrifugation for 30sec at 3000rpms and the supernatant was collected and transferred to a new tube. To reverse crosslinking, each sample received 30uL of 5M NaCl and incubated at 65°C overnight. The following day, ChIP DNA was purified using a PCR clean up kit (Qiagen) and concentrated into 35uL of water. Enrichment of immunoprecipitated DNA was quantified using the Biorad 2x SYBR Green Master mix, 2ul of ChIP DNA per reaction and primers at a final concentration of 250nM. For a list of primer sequences, see supplemental table 2.

### Stacked oocyte scoring and rescue

*heri-1* mutant or control animals were crossed to N2 males and maintained as heterozygous for 1 to 2 generations. Homozygous *heri-1* mutant animals were isolated and stacked oocytes were quantified in multiple siblings the next generation. This strategy was adopted to minimize the chance that transgenerational epigenetic effects could accruing in *heri-1* lines prior to analysis. To score stacked oocyte defects, L4 animals were placed in OP50 food at 20°C for 24hrs, then placed in M9 buffer with 0.1% Sodium Azide and mounted onto 2% agarose pads. For each biological replicate, 50 worms were scored. To ask if mating with wild-type males could rescue stacked oocyte defects, we singled L4 stage *heri-1(-); pie-1::gfp::h2b* animals and allowed these animals grow at 20° for 24hrs. Animals with stacked oocytes in both gonad arms were identified with a dissecting microscope and split into two groups. Group #1 were not crossed to wild-type males (No cross). Group #2 worms were mated with three wild-type males (Cross). Approximately 48hrs later, males were removed and animals were allowed to grow at 20°C for another 24 hrs. The number of progeny for each worm in each group was counted.

### Immunofluorescence and imaging

For HERI-1::3xFLAG imaging, gonads were dissected and fixed as follows: ~20 gravid hermaphrodites where picked onto a 20uL drop of M9 in a coverslip. Gonads were dissected using a 25G ⅝ needle by cutting worms in half near vulva. Coverslip containing dissected gonads was placed in a Gold Color Frost Plus slide (9951GLPLUS, ThermoFisher) and immediately placed on a block of dry ice for 10 mins. Using a razor blade, the coverslip was popped off the sample, then quickly place in -20°C cold MeOH for 10 min. The slide was allowed to air dry for 1-2 min at room temperature. Excess ethanol was removed with a kimwipe. Samples were blocked with 500uL of 0.5%BSA in 1xPBS at room temperature. The M2 anti-flag antibody (F1804 Sigma) was diluted 1:100 in 0.5%BSA in 1xPBS and 50uL was added to the sample. Primary incubation was for overnight at 4°C in a humidified chamber. Slides were then washed 4x with 150uL of 0.5%BSA in 1xPBS with 6-7 mins for each wash. An Alexa Fluor 488 goat anti-mouse secondary antibody (A10667, Molecular Probes) was diluted 1:250 in 0.5%BSA in 1xPBS and 50uL were added to the sample. Secondary incubation was for 2 hrs at room temperature. Slides were again washed 4x with 150uL of 0.5%BSA in 1xPBS with 6-7mins each wash. After final wash, 15uL of Vectashiled with DAPI (H-1200, Vector Laboratories) was added to the slide and a new coverslip was carefully placed over the sample. Images were taken with our One Zeiss Observer.Z1 inverted confocal microscope.

## Supplemental data

**Figure S1.**
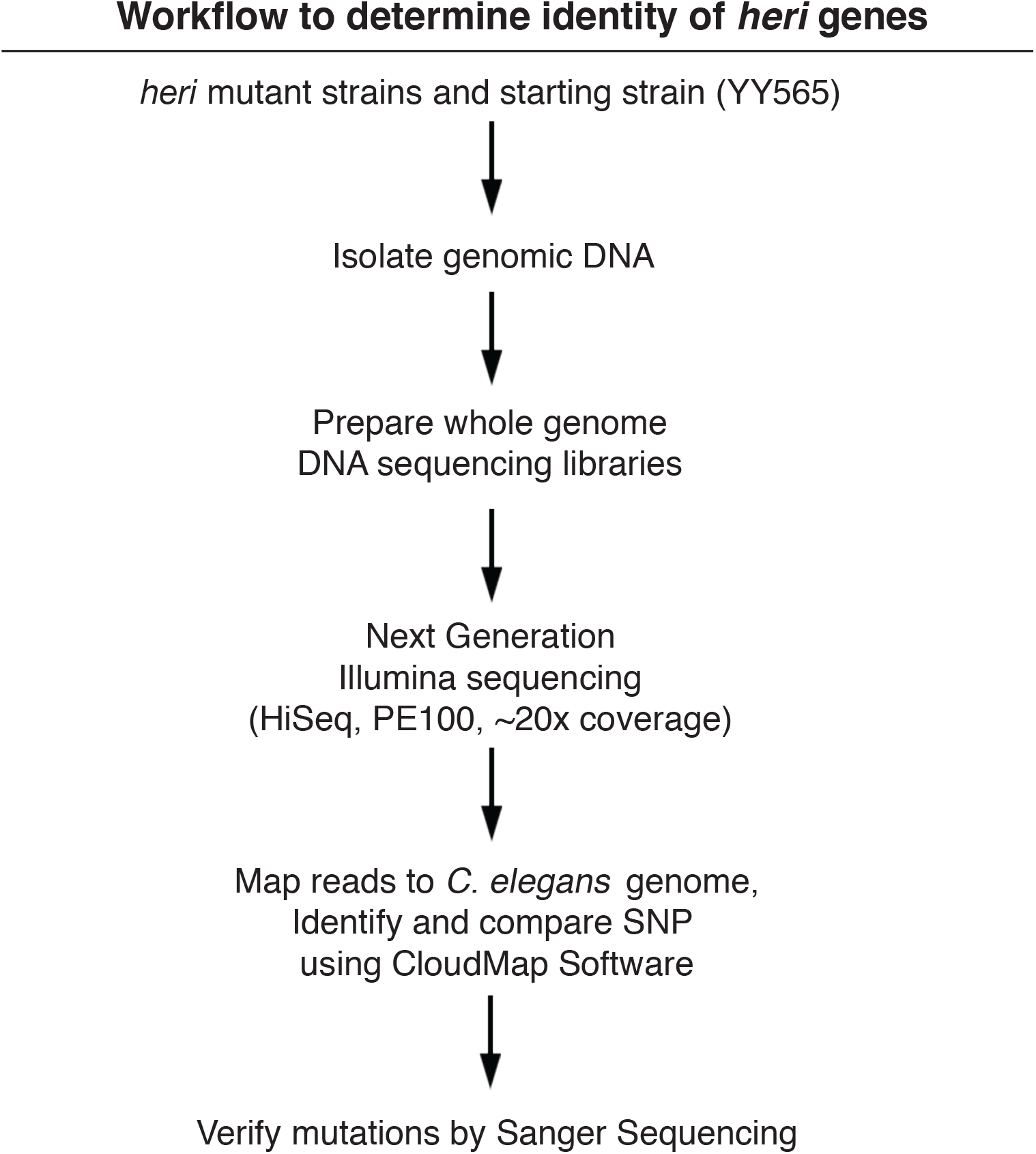
Identifying Heri genes. Workflow used to identify *heri-1*. CloudMap software is described in (Minevich et al. 2012).

**Figure S2.**
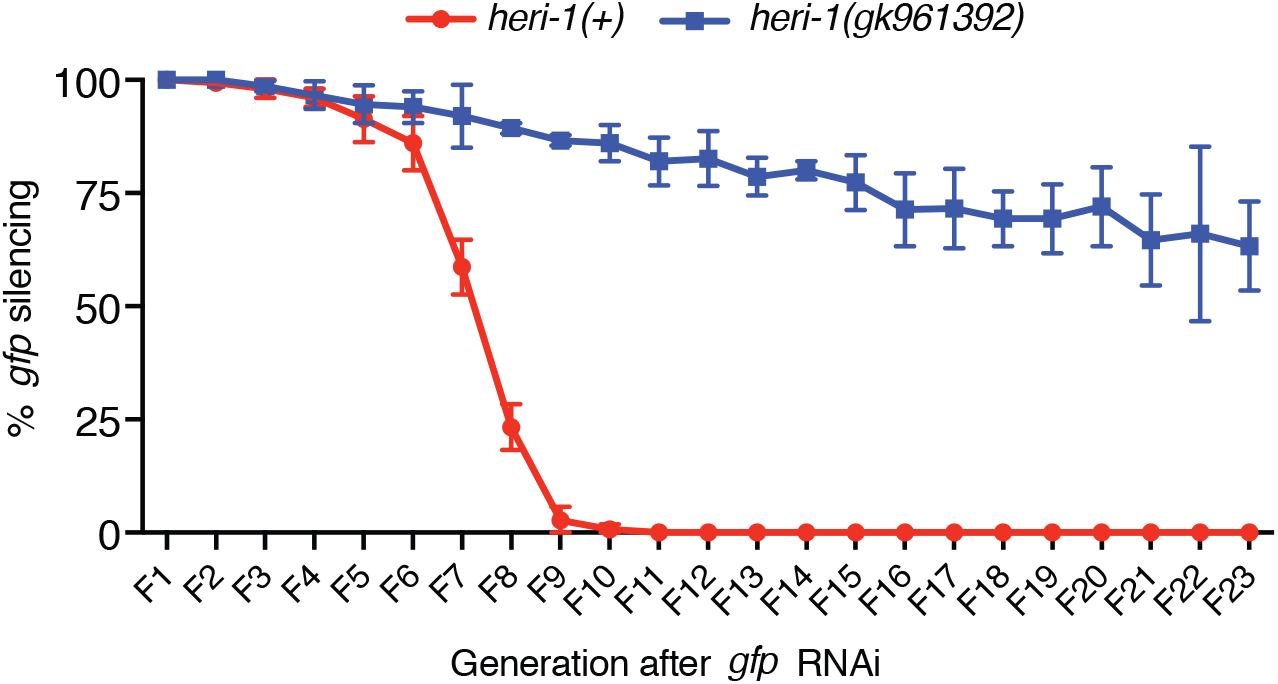
*heri-1(gk961392*) enhances TEI. Animals of indicated genotypes that expressed a *pie-1::h2b::gfp* transgene were treated with *gfp* RNAi (see methods). Data points represent 3 biological replicates in which 50 animals were scored for GFP expression. Experiment was performed blind and error bars represent standard deviations of the mean.

**Figure S3.**
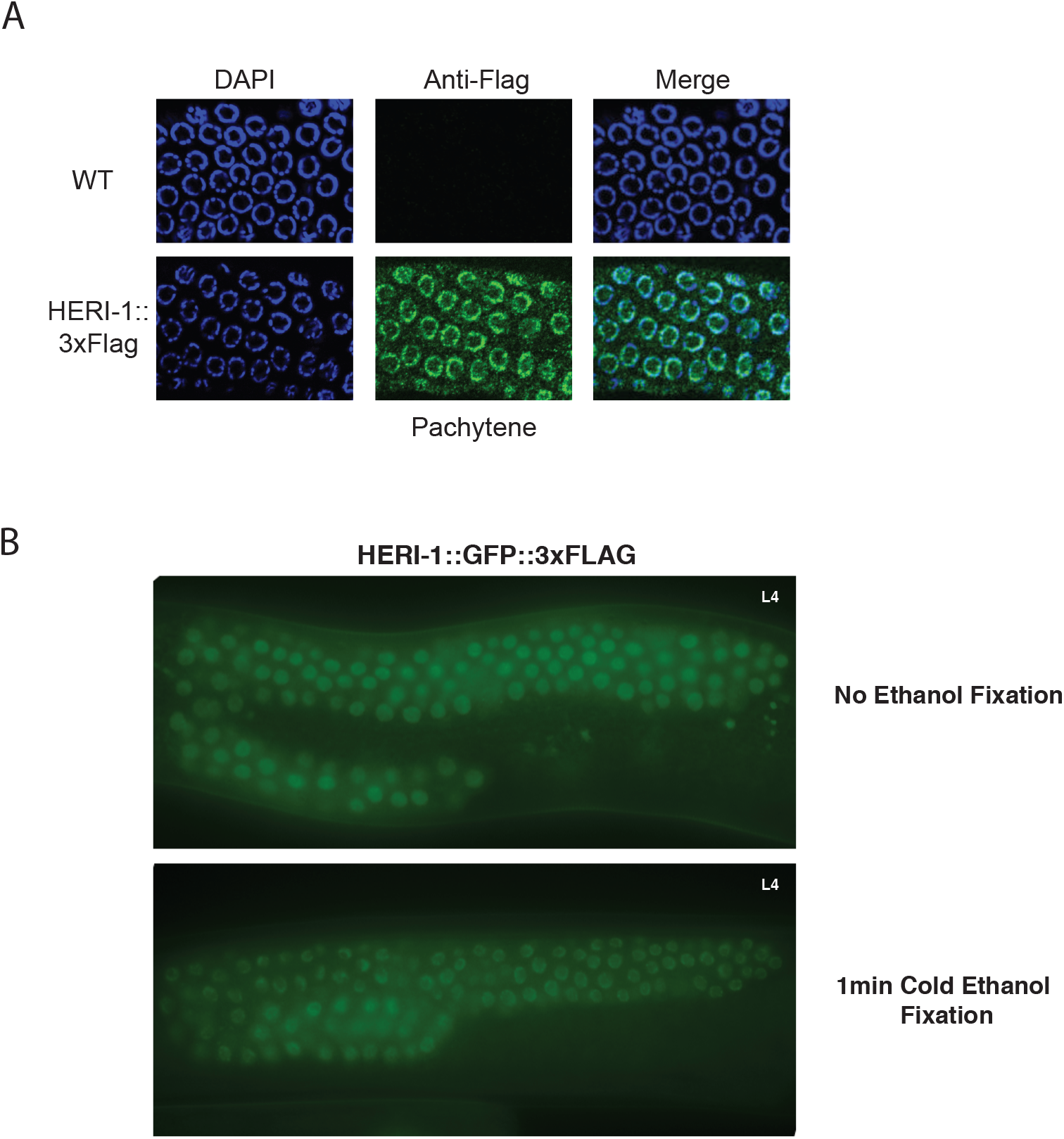
HERI-1 localizes to nuclei. **A)** Micrographs show immunofluorescent (IF) images of fixed adult pachytene germ cell nuclei in wild type and HERI-1::3xFLAG detected with α-FLAG antibodies. DAPI staining (DNA) is shown in blue. Note that the subnuclear distribution of HERI-1 appears more peripheral in these micrographs that when we visualized HERI-1::GFP in the main text (Fig. 3A). The data in (**B)** hint at why this might be. **B)** Micrographs of larval stage 4 (L4) germlines without ethanol fixation (top panel) or with -20°C ethanol fixation for 1 min (bottom panel). During ethanol fixation, germlines of L4 animals were dissected onto a subbed slide, placed on a cover slip, and snap frozen on dry ice for 10 mins. Cover slips were removed and the gonads were fixed in -20°C ethanol in a coplin jar for 1 min. Slides were allowed to air dry for 5 mins and excess ethanol was removed and gonads were imaged under our inverted microscope. This fixation process seems to alter the sub-nuclear distribution of HERI-1 in a way that makes HERI-1 appear to colocalize with chromatin. Notice the localization of HERI-1::GFP::3xFlag closer to the nuclear periphery after fixation.

**Figure S4.**
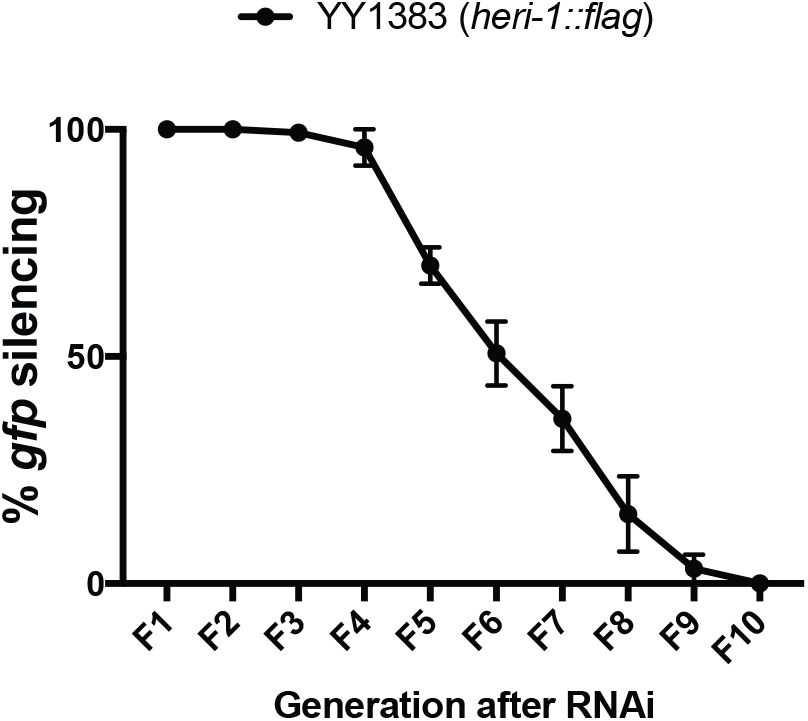
HERI-1::3xFLAG is functional for RNAi inheritance. *heri-1::flag* animals that express the *pie-1::h2b::gfp* transgene were treated with *gfp* RNAi (see methods) and the percentage of animals expressing GFP across generations is indicated. Data points represent 3 biological replicates in which 50 animals were scored for GFP expression. Experiment was performed blind and error bars represent standard deviations of the mean. Because *gfp* inheritance silencing looks similar to what we always see in wild type animals (compare Figure S2 and Figure S3), we conclude that HERI-1::FLAG is functional. Note, we have not tested HERI-1::GFP::FLAG animals in this assay because these animals harbor two *gfp* loci.

**Figure S5.**
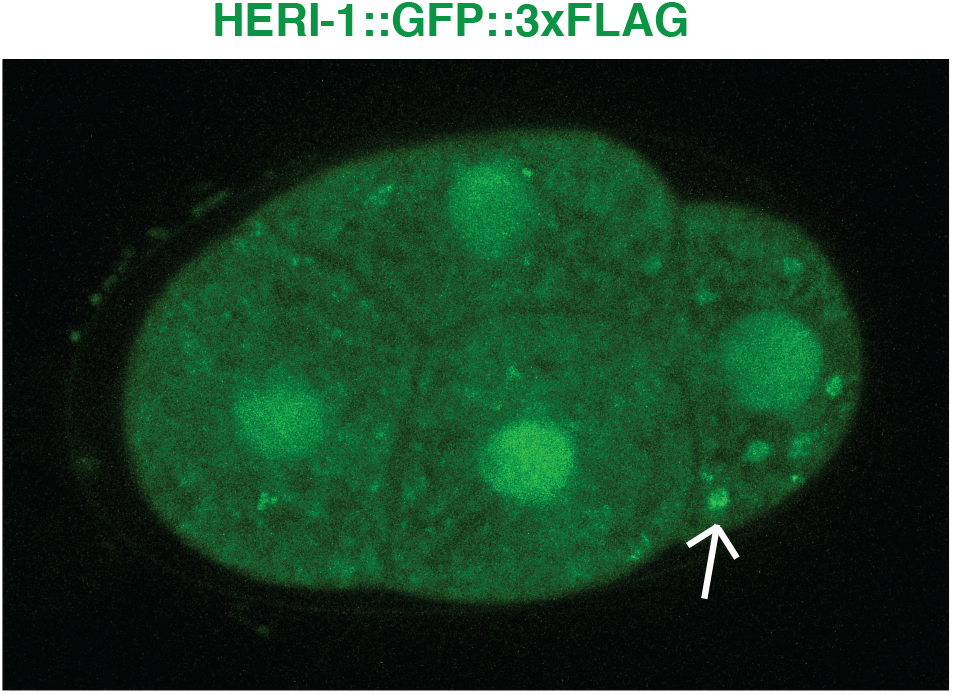
HERI-1 is expressed in somatic and germline blastomeres during early development. Micrograph of an 4 cell embryo expressing HERI-1::GFP::3xFLAG. HERI-1 is present in all four nuclei. In the P2 cells, some cytoplasmic localization is seen that may indicate that some HERI-1 is present of P-granules in this cell (white arrow).

**Figure S6.**
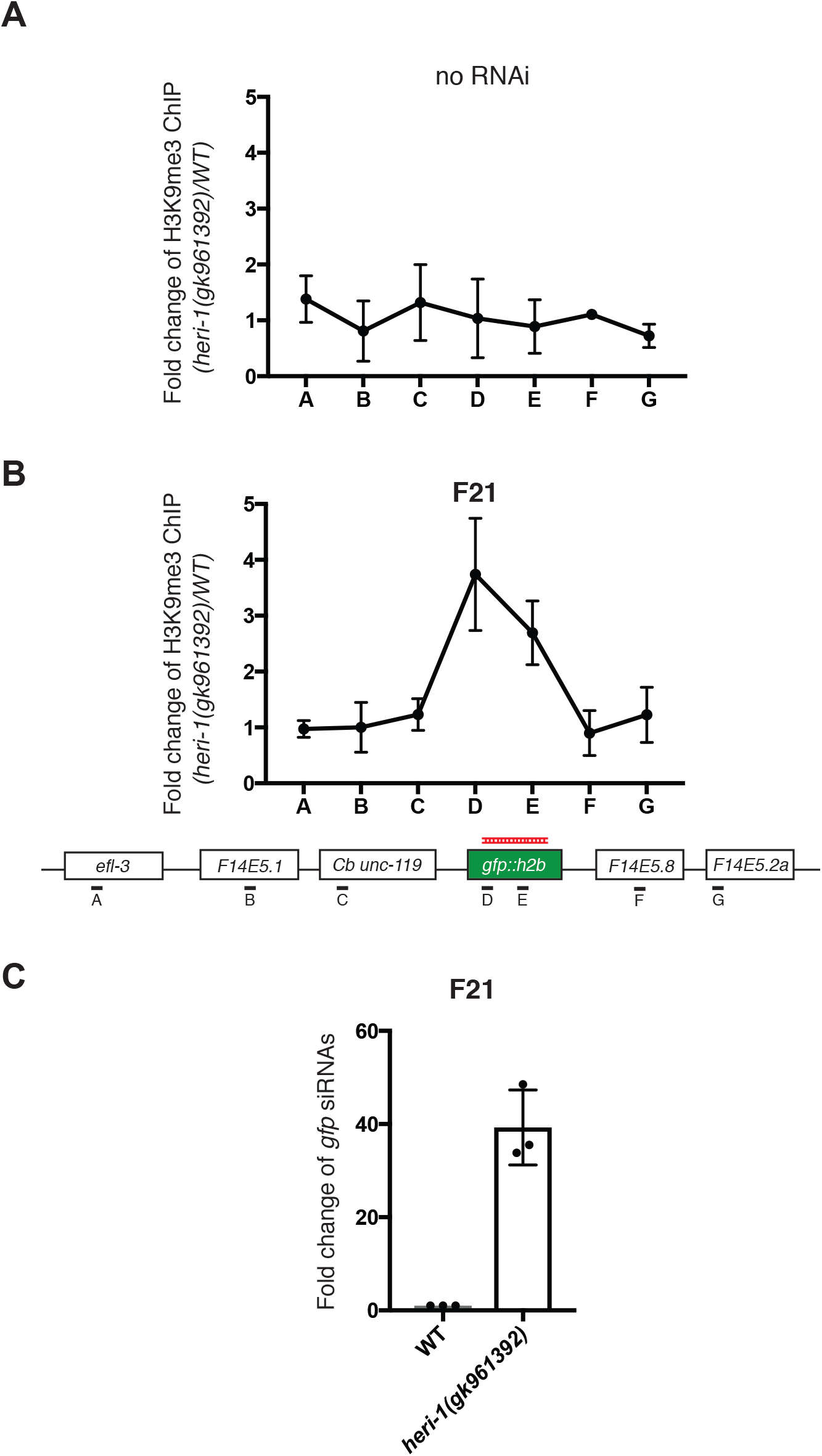
*heri-1(-)* animals exhibit elevated levels of H3K9me3 and siRNAs twenty generations after RNAi. **(A/B)** H3K9me3 ChIP-qPCR on *heri-1(+) heri-1(-)* animals. Data is expressed as a ratio of H3K9me3 ChIP signals detected in *heri-1(+*) over *heri-1(-)* mutant animals before *gfp* RNAi (bottom panel) and twenty generations after *gfp* RNAi (bottom panel). Data is from 3 biological replicates and error bars are standard deviations of the mean. **C)** Total RNA isolated from animals of the indicated genotypes was used for Custom Taqman assays (Burton et al, 2011) to detect *gfp* siRNAs twenty generations after *gfp* RNAi treatment. Signal from WT (starting strain) animals is defined as one. Data is from 3 biological replicates and error bars are standard deviations.

**Figure S7.**
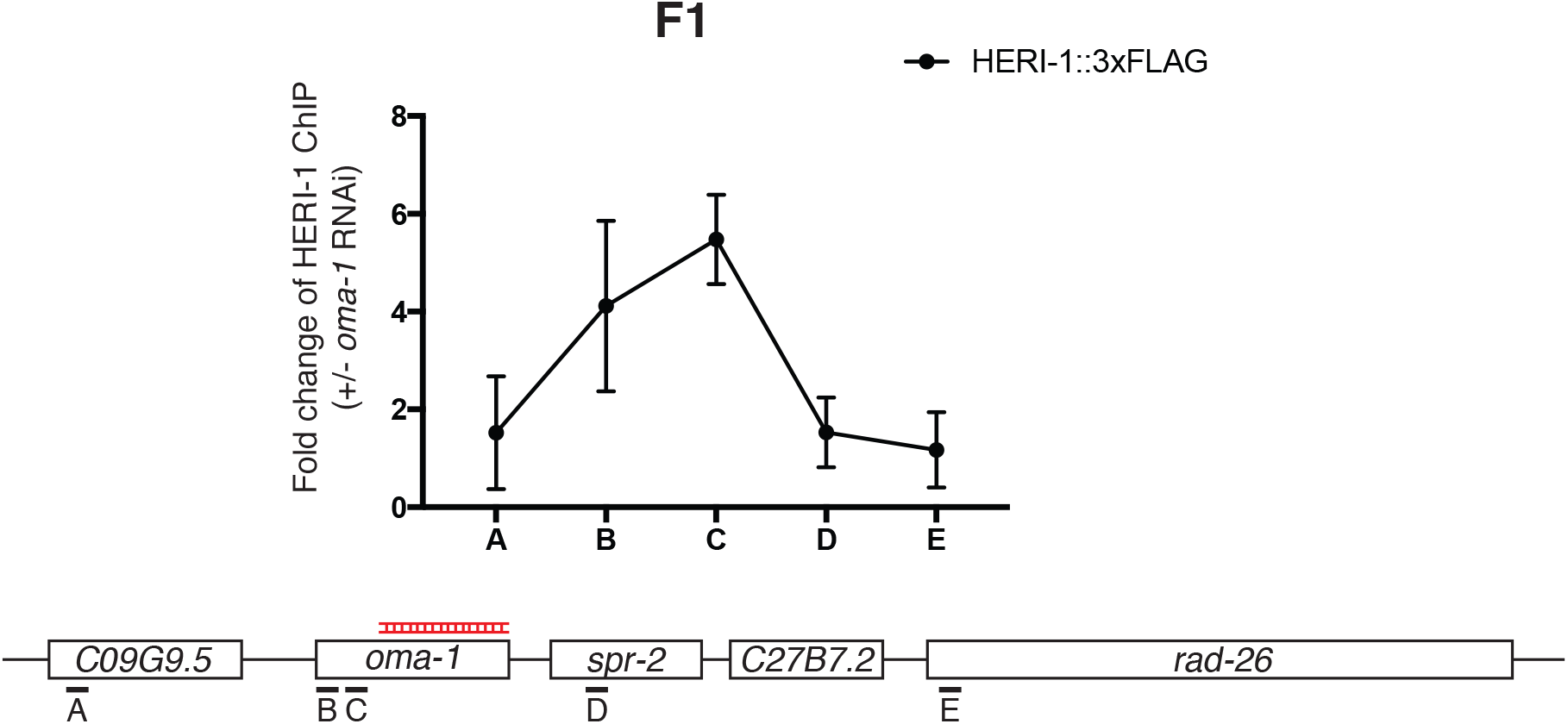
HERI-1::3xFLAG binds to chromatin in response to RNAi. FLAG ChIP-qPCR on progeny of HERI-1::3xFLAG animals treated +/-*oma-1* RNAi. Data is expressed as a ratio of FLAG ChIP signals in animals treated with *oma-1* RNAi over non-RNAi control animals. Data is from 3 biological replicates and error bars are standard deviations of the mean. Pattern is similar to what we observed with HERI-1::GFP in Fig. 5.

**Figure S8.**
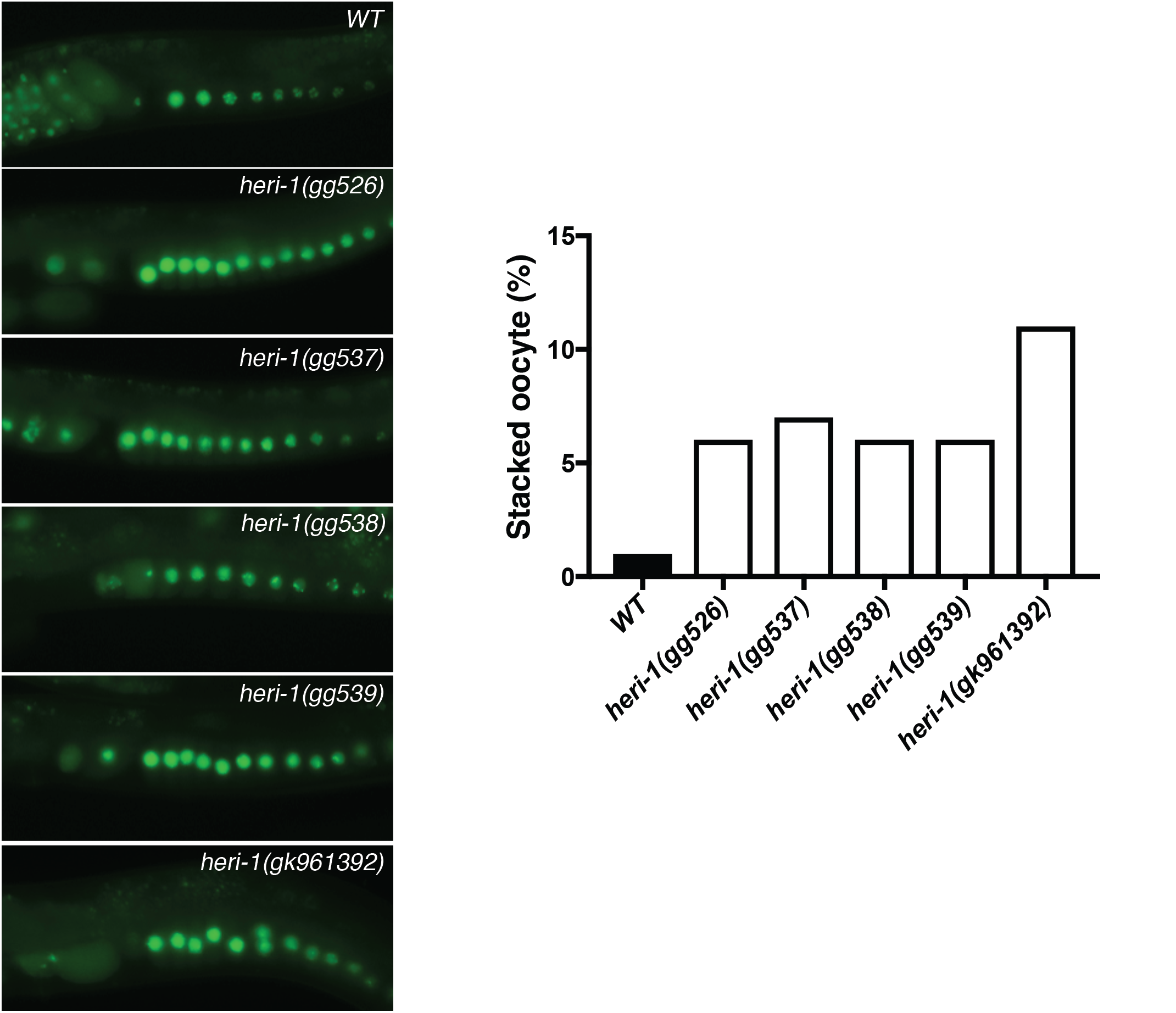
*heri-1* mutant animals show a “stacked oocyte” phenotype. Micrographs of stacked oocyte defects in five different *heri-1* mutants strains. *gg526, gg537, gg538* and *gg539* were from Heri screen. *gk961392* was from the million mutation project. All animals are expressing a *pie-1::gfp::h2b* reporter gene. Quantification (50 animals scored/strains) is shown on the right.

**Figure S9.**
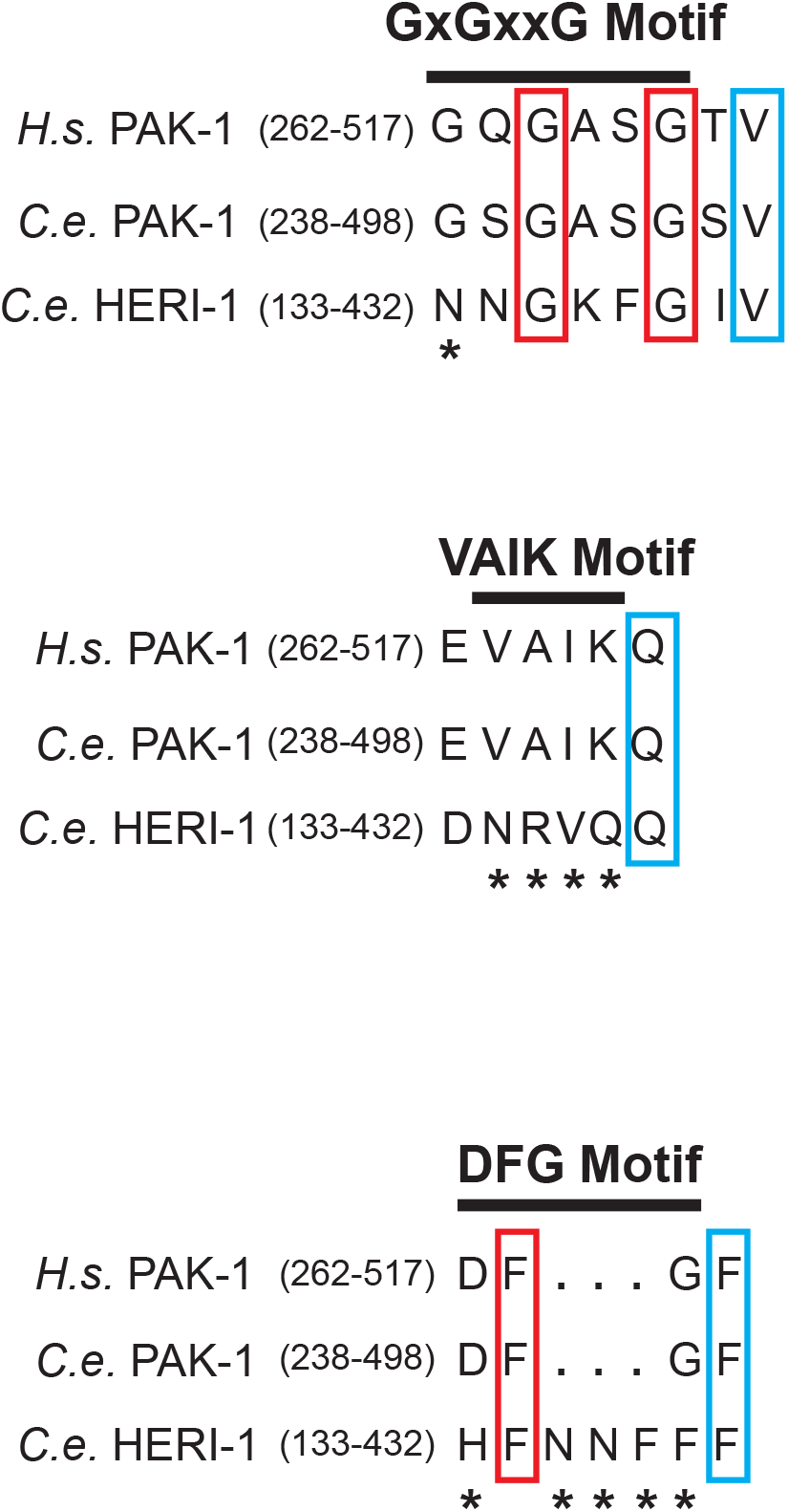
HERI-1’s kinase motifs are degenerate. Sequence alignments of three motifs (GxGxxG VAIK and DFG), which contribute to kinase activity in most protein kinases, from human PAK-1, C. *elegans* PAK-1 and HERI-1. Motif location is indicated by the black bar. Amino acids in red rectangles indicate conservation within the motif, while amino acids in blue rectangles indicate conservation outside the motif. Asterisks indicate lack of conservation.

**Figure S10.**
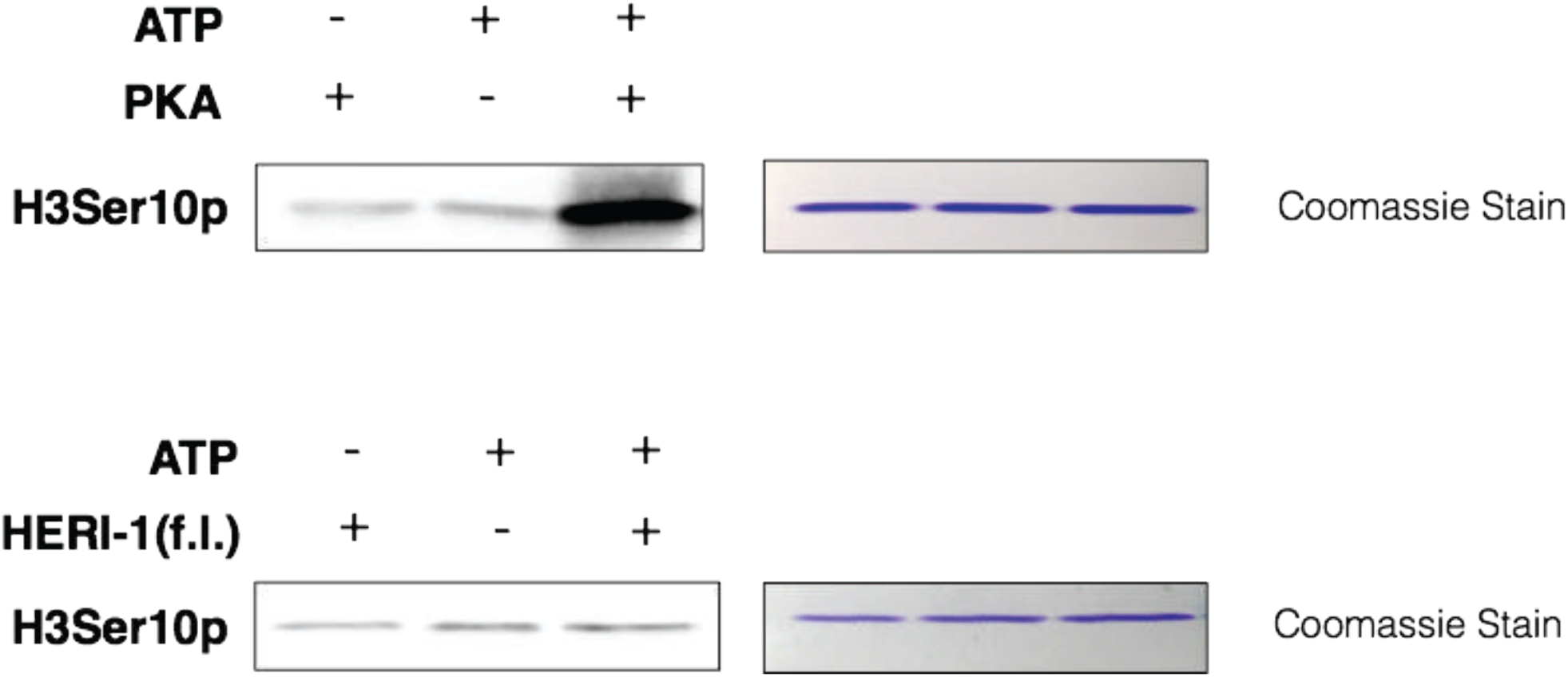
No kinase activity detected with recombinant HERI-1 protein *in vitro*. 1ug of purified Histone H3 was incubated with +/-ATP and +/-cAMP-dependent protein kinase (PKA) (NEB, cat. no P6000S) (top panel), or with full length HERI-1 recombinant protein (bottom panel). Kinase reactions were performed in kinase assay buffer (200mM final ATP and 1x Kinase Buffer (NEB) for 1hr at 37° in a final volume of 15uL). After incubation, 15uL of 2x Loading buffer were added to each reaction. Incubated at 95° for 5 min, and resolved in a 7.5% Polyacrylamide gel. Detection of H3Ser10p was determined by using the anti-H3Ser10p antibody from Millipore (clone CMA312) at a dilution of 1:1000. Purified Histone H3 samples stained with coomassie blue are shown as loading controls. Full length (f.l).

**Figure S11.**
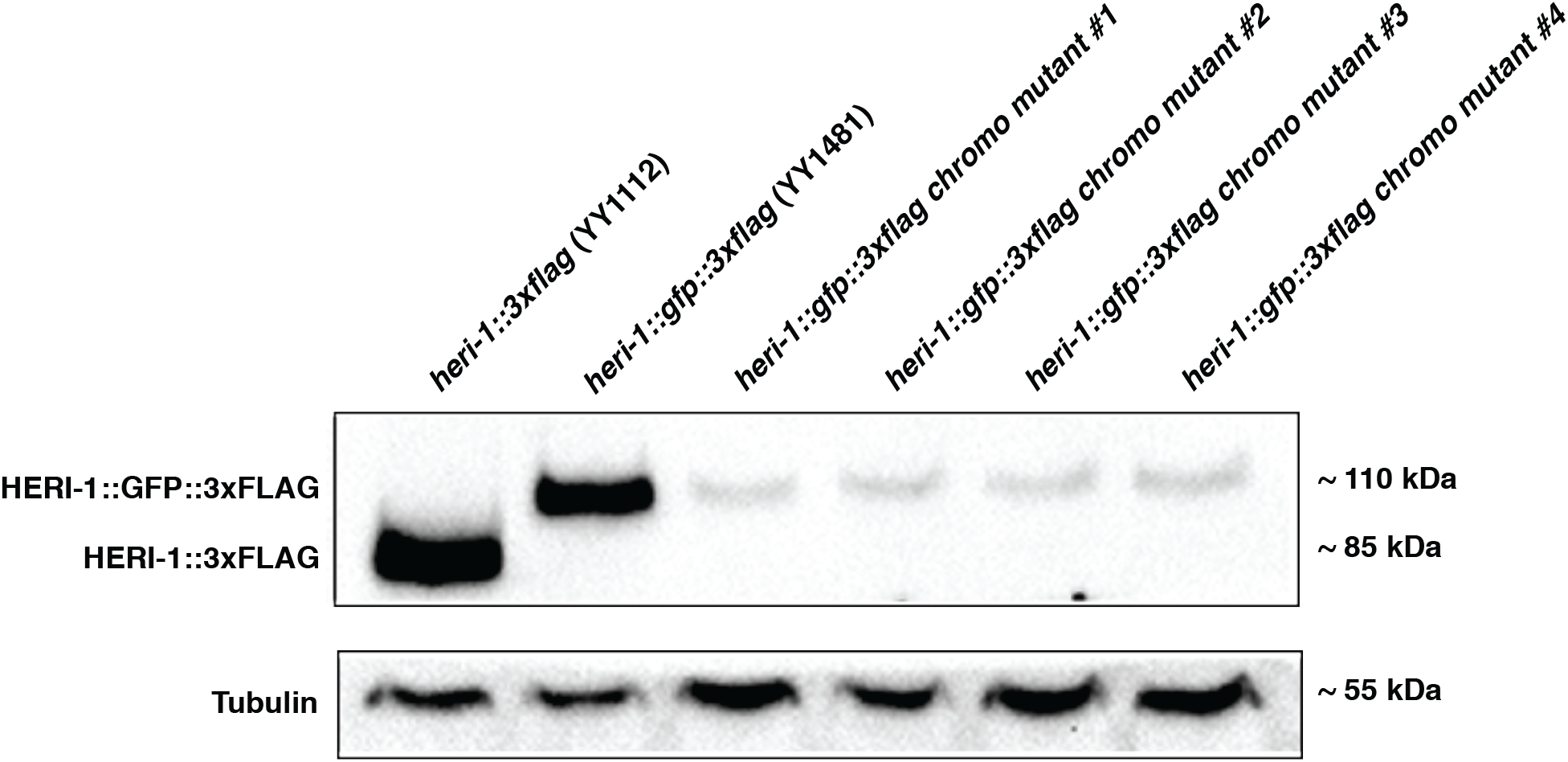
HERI-1 chromodomain mutant protein (*gg526*) is unstable. *gg526* is a *heri-1* mutation identified by our screen that mutates the HERI-1 chromodomain (V60D, see Fig. 2). Western blot analysis comparing levels of HERI-1::3xFlag, HER-1::GFP::3xFlag, and HERI-1(V60D)::GFP::3xFLAG is shown. α-Flag antibody (Sigma, M2) was used to detect proteins and a α-tubulin antibody was used for loading control. Shown are 4 lines (mutant 1-4) of HERI-1(V60D)::GFP::3xFLAG animals produced by CRISPR-based *gfp::3xflag* tagging of *heri-1(gg526*).

---

**Supplemental Table 1.** Strains used in this study

| **<u>Name</u>** | **<u>Genotype</u>** | **<u>Source</u>** |
| --- | --- | --- |
| N2 | *Wild Type* | CGC |
| YY565 | *oma-1(zu405) IV; mjlS31(pie-1::gfp::h2b) II* | This study |
| YY640 | *hrde-1(tm1200) III; mjlS31 II* | Buckley et.al. |
| YY942 | *heri-1(gg539) II; oma-1(zu405) IV; mjlS31 II* | This study |
| YY941 | *heri-1(gg538) II; oma-1(zu405) IV; mjlS31 II* | This study |
| YY940 | *heri-1(gg537) II; oma-1(zu405) IV; mjlS31 II* | This study |
| YY939 | *heri-1(gg526) II; oma-1(zu405) IV; mjlS31 II* | This study |
| YY1032 | *heri-1(gk961392) II ox=4* | This study |
| YY1112 | *heri-1::3xflag II* | This study |
| YY1218 | *cec-9(gk961392) II;mjlS31 II* | This study |
| YY1224 | *heri-1::3xflag II; hrde-1(tm1200) III* | This study |
| YY1228 | *heri-1::3xflag II; hrde-3(gg546) I* | This study |
| YY1481 | *heri-1::gfp::3xflag II* | This study |
| YY1500 | *heri-1::gfp II;mcherry::h2b* | This study |
| YY1642 | *heri-1(gk961392) II; hrde-1(tm1200) III; mjlS31 II* | This study |
| VC40608 | *cec-9(gk961392) II* | Million Mutation Project |
| SX461 | *mjlS31(pie-1::gfp::h2b) II* | Bagijn et.al. |

